# Analyzing ER stress response in ALS patient derived motor neurons identifies druggable neuroprotective targets

**DOI:** 10.1101/2023.11.16.567279

**Authors:** Michelle E. Watts, Richard M. Giadone, Alban Ordureau, Kristina M. Holton, J. Wade Harper, Lee L. Rubin

## Abstract

Amyotrophic lateral sclerosis (ALS) is a degenerative motor neuron (MN) disease with severely limited treatment options. Identification of effective treatments has been limited in part by the lack of predictive animal models for complex age-related human disorders. Here, we utilized pharmacologic ER stressors to exacerbate underlying sensitivities conferred by ALS patient genetics in induced pluripotent stem cell (iPSC)-derived motor neurons (MNs). In doing so, we found that thapsigargin and tunicamycin exposure recapitulated ALS-associated degeneration, and that we could rescue this degeneration via MAP4K4 inhibition (MAP4K4i). We subsequently identified mechanisms underlying MAP4K4i-mediated protection by performing phosphoproteomics on iPSC-derived MNs treated with ER stressors -/+ MAP4K4i. Through these analyses, we found JNK, PKC, and BRAF to be differentially modulated in MAP4K4i-protected MNs, and that inhibitors to these proteins could also rescue MN toxicity. Collectively, this study highlights the value of utilizing ER stressors in ALS patient MNs to identify novel druggable targets.

## 1 Introduction

Amyotrophic lateral sclerosis (ALS) is a fatal neurodegenerative disorder characterized by selective death of motor neurons (MNs) (Brown and Al-Chalabi 2017). The loss of this cell type in the brain and spinal cord manifests in symptoms of muscle weakness and atrophy, which progress rapidly to paralysis and respiratory failure approximately 1-5 years after diagnosis (Hardiman et al. 2017). Currently, there are only 4 FDA approved treatments for the disease (riluzole, edaravone, and the recently approved relyvrio and tofersen). However, none of these extend life expectancy beyond several months nor improve muscle function in all patient cohorts (Jaiswal and M.K. 2019), and only tofersen acts via a widely accepted disease-associated mechanism. Developing efficacious therapeutics for ALS has been particularly challenging in part due to the inaccessibility of human tissue for study. Moreover, animal models fail to recapitulate the variable genetic drivers observed in patients, including coding and non-coding elements, thereby limiting their effectiveness and therapeutic translatability (Petrov et al. 2017).

To overcome this limitation, several studies have employed patient-specific induced pluripotent stem cell (iPSC)-based culture systems to understand ALS pathogenesis as well as evaluate the ability of pre-clinical therapeutic candidates to attenuate disease phenotypes (Dimos et al. 2008; Fujimori et al. 2018; Wainger et al. 2014). iPSC lines have been generated from individuals with a variety of mutations in genetic drivers of the disease (e.g., FUS, C9ORF72, TDP43, SOD1) as well as more common sporadic forms (Boulting et al. 2011; Burkhardt et al. 2013; Ichiyanagi et al. 2016; Kiskinis et al. 2014; Sareen et al. 2013). Through these models, the field has recapitulated a number of disease-relevant phenotypes in iPSC-derived neuronal cell types, including: altered neurite morphology (Chen et al. 2014; Egawa et al. 2012; Fujimori et al. 2018), disease-specific transcriptional signatures (Workman et al. 2023), increased excitotoxicity (Shi et al. 2018; Wainger et al. 2014), accumulation of intracellular aggregates (Egawa et al. 2012; Hung et al. 2023; Naumann et al. 2018; Shi et al. 2019; Sun et al. 2018), mRNA mis-processing (Melamed et al. 2019), increased apoptotic activity (Abo-Rady et al. 2020; Kiskinis et al. 2014; Naujock et al. 2016; Wu, Watts, and Rubin 2019), and increased sensitivity to stressors (Shi et al. 2018; Yang et al. 2013; Zhang et al. 2013). Utilizing these models, it may be possible to identify convergent mechanisms driving MN death in ALS patients, perhaps enabling the development of broadly efficacious therapeutics.

In addition to patient genetics, aging is a significant risk factor for the development of ALS, as it is for many other prominent neurodegenerative diseases (Hardiman et al. 2017). With age, patients experience loss of protein homeostasis (proteostasis) networks resulting in accumulation of misfolded proteins and ER stress (Hipp, Kasturi, and Hartl 2019). Accumulation of aggregated proteins in the ER lumen can lead to activation of several signaling pathways, including the unfolded protein response (UPR) (Matus, Glimcher, and Hetz 2011). Specifically, the ER chaperone BiP dissociates from three transmembrane receptors (ATF6, IRE1, and PERK) and binds to misfolded proteins. This initial dissociation of BiP leads to activation of downstream signaling events, including proteolytic cleavage of ATF6 in the Golgi, phosphorylation of ATF4 downstream of PERK and eIF2α phosphorylation, and splicing of XBP1 mRNA (downstream of IRE1 activation). Ultimately, these events lead to induction of transcription factors that translocate into the nucleus and activate downstream transcriptional networks with adaptive/cytoprotective or pro-apoptotic outputs including: upregulation of chaperone genes downstream of ATF6, ER associated degradation (ERAD) components via XBP1, and pro-apoptotic machinery via PERK (Hetz, Zhang, and Kaufman 2020). MNs, with characteristically long axonal projections exhibit increased demand on ER-Golgi secretory pathways, ultimately resulting in high levels of basal ER stress. Further, in the case of ALS MNs, several studies demonstrate pathogenic aggregation of disease-associated proteins (e.g., FUS, TDP-43, and SOD1) contributes to increased UPR activity. Coupled with increased basal activation of ER stress, this leads to increased sensitivity to apoptotic signaling, and ultimately death of vulnerable MN subpopulations, implicating proteostasis dysfunction in the pathobiology of ALS (Rozas et al. 2017; Ruegsegger and Saxena 2016; Saxena, Cabuy, and Caroni 2009; Webster et al. 2017).

A current challenge in the field of iPSC-based modeling of neurodegenerative diseases such as ALS has been recapitulating the age-related decline of fundamental processes such as proteostasis. This is especially true as the reprogramming process used to generate iPSCs eliminates epigenetic changes associated with aging. Additionally, one hallmark of ALS often overlooked in cell culture models includes differential MN vulnerability, where upper MNs in the motor cortex of the brain and lower MNs in the brainstem and spinal cord are disproportionally damaged in patients relative to other CNS cell types (Ferraiuolo et al. 2011; Ragagnin et al. 2019). Studies utilizing stem cell-based differentiation platforms have predominately focused on minimizing heterogeneity in their cultures, potentially overlooking important differences in cell type-specific responses to proteostatic perturbations.

To address these challenges and better recapitulate the MN-specific vulnerability observed in patients with ALS, we employed an established MN differentiation protocol producing cells of lower motor column (LMC) identity, including: MNs, ventral spinal interneurons, and a small number of astroglia (Maury et al. 2014; Neel et al. 2023). We then exposed these cells to chemical inducers of ER stress to exacerbate underlying ER stress signaling. We hypothesized that stressors preferentially affecting the MN cell types within these heterogenous cultures may reflect selective damage of MNs within the diverse spinal cord niche. Through these efforts, we observed that 2 mechanistically distinct ER stressors, thapsigargin (a SERCA inhibitor) and tunicamycin (an N-linked glycosylation inhibitor) synchronously recapitulated the activation of early disease-associated UPR signaling events, along with late-stage MN-specific degenerative phenotypes in both healthy and ALS patient MNs. We then validated the fidelity of this approach in identifying neuroprotective drugs by reproducing the ability of several pharmacological MAP4K4 inhibitors, originally identified by our group, to rescue this ALS MN toxicity. To discover additional neuroprotective effectors downstream of MAP4K4 MN protection, we subsequently performed phosphoproteomics on patient derived MNs after exposure to thapsigargin, with or without protection by MAP4K4 inhibition. In doing so, we identified PKC and BRAF as signaling components downstream of MAP4K4 inhibition, and further showed that commercial inhibitors of these targets also exhibited neuroprotective effects. Collectively, these data demonstrate the effectiveness of utilizing chemical ER stress to exacerbate ALS disease phenotypes in patient-derived cells and that synergizing this platform with proteomics assays allows for identification of druggable targets for neurodegenerative diseases.

## 2 Results

### 2.1 ER stress preferentially induced death in MN, but not non-MN, ALS iPSC-derived cell populations

Many studies have implicated ER stress as an underlying mechanism of MN death in ALS, demonstrating changes in solubility and localization of intracellular aggregates upon treatment with ER stress-inducing compounds such as thapsigargin, tunicamycin, and MG132 (Bhinge et al. 2017; Medinas et al. 2018; Walker et al. 2013); however, few have investigated toxicity or unbiased proteomic/phosphoproteomic changes upon perturbation. To investigate the extent to which proteostatic stress initiates preferential MN degeneration in human cells, we first evaluated MN cultures derived from a non-diseased, healthy control 1016A hiPSC line. hiPSCs were differentiated into MNs following an established 15-day embryoid body (EB)-based protocol that recapitulates neurectoderm induction with dual SMAD inhibition, caudalization with retinoic acid (RA), ventralization with Sonic Hedgehog pathway activation, and MN maturation with the neurotrophic factors (BDNF, GDNF), and γ-secretase inhibitor DAPT (Figure 1A)(Maury et al. 2014). Following this protocol, ∼80% of cells were neurons (based on the expression of βIII-Tubulin; TUJ1+), ∼30% of which expressed the mature MN marker Isl1/2 (Isl1/2+, TUJ1+) (Figure 1B-C). ISL1/2 and TUJ1 double-positive cells were considered MNs, while the remaining populations were considered non-MN cells (Figure 1B)(Neel et al. 2023).

**Figure 1.**
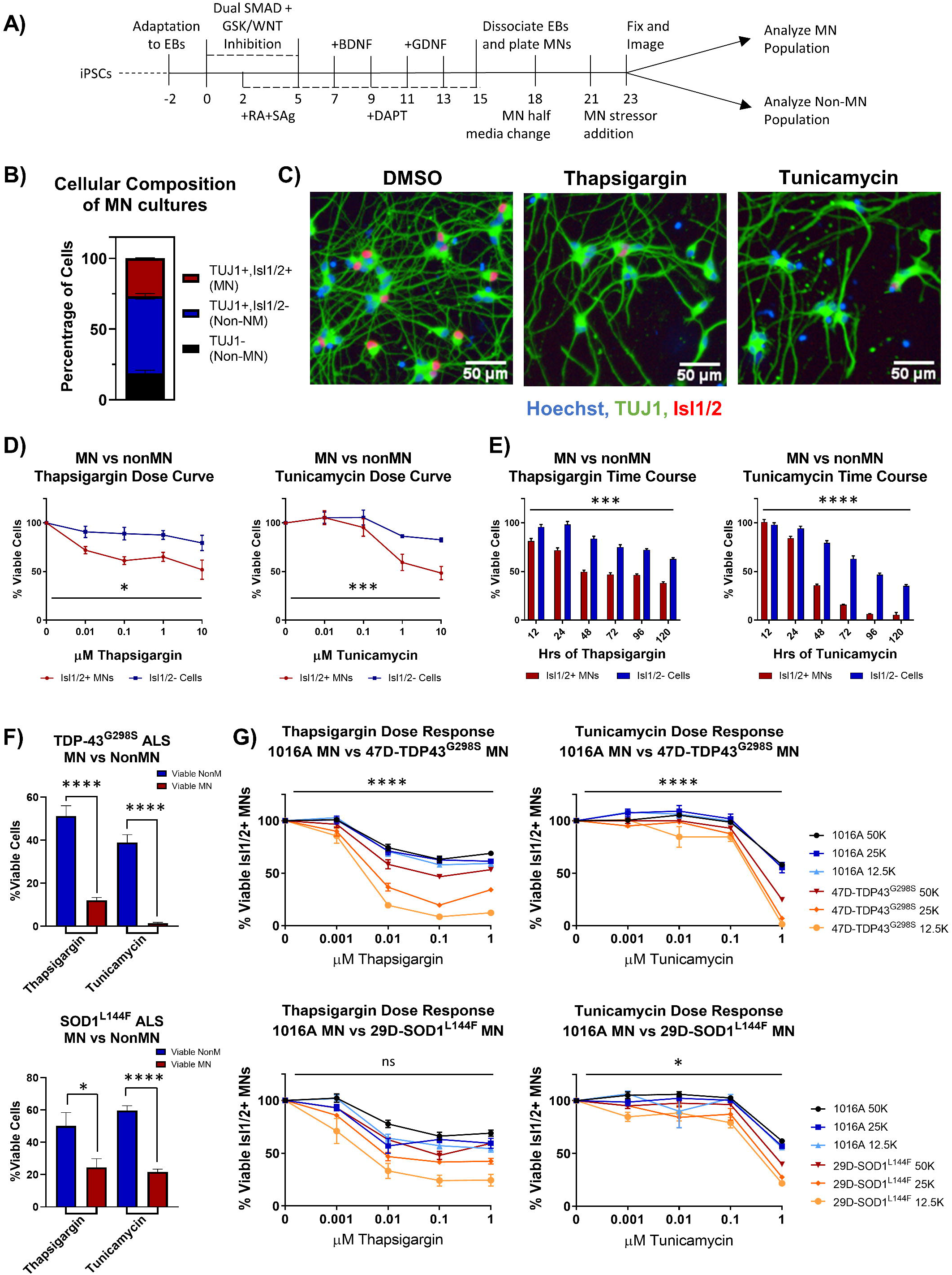
ER stressors induce preferential MN death. (A) Schematic of MN differentiation protocol and stressor assay. (B) Quantification of cell types in patient derived MN cultures. MN populations include Isl1/2+, TUJ1+ immunostained cells, non-MN populations include Isl1/2-, TUJ1+ neuronal populations and TUJ1-nonneuronal cells. Nb=4, nt=6. (C) Representative immunofluorescent images of patient derived MN cultures in control (DMSO) or ER stress (1μM compound, 48hrs) conditions. (D) Quantification of MN and non-MN viability 48hrs after treatment with increasing concentrations of ER stressors. Nb>3, nt>2, two-way ANOVA; Thapsigargin- p = 0.015, Tunicamycin- p = 0.000127. (E) Quantification of MN and non-MN viability after treatment with 1μM ER stressors for various lengths of time. Nb=3, nt=6, two-way ANOVA; Thapsigargin- p = 0.000215, Tunicamycin- p = 1.64×10^-33^. (F) Comparison of viable non-MNs and viable MNs after TDP-43^G298S^ mutant or SOD1^L144F^ mutant ALS patient derived MN cultures were treated with 48hrs 1μM ER stressors. Nb=3, nt >2, 2 tailed unpaired students t-test; 47D-Thapsigargin- p = 1.526×10^-5^, 47D-Tunicamycin- p =1.450×10^-6^, 29D-Thapsigargin- p = 0.02698, 29D-Tunicamycin- p = 6.07742×10^-7^. (G) Quantification of viable MNs from a healthy patient line (1016A) and ALS patient lines (47D-TDP-43^G298S^ or 29D-SOD1^L144F^) after 48hrs of ER stressor exposure. 4 different concentrations of each stressor were tested, in 3 different densities of each line. Nb=3, nt=2, three-way ANOVA comparing interaction of cell density and stressor dosage between ALS and control; 47D-Thapsigargin- p = 5.81×10^-11^, 47D-Tunicamycin- p = 1.37×10^-17^, 29D-Thapsigargin- p = 0.515, 29D-Tunicamycin- p= 0.016. Biological replicate experiments denoted as Nb, each with technical replicate experiments nt. Data are mean value +/- SEM.

Since preferential death of MNs in ALS occurs in the spinal cord, comprised of diverse cell types, we then compared the response of MN and non-MN populations in our cultures to challenge with proteotoxic stressors. In doing so, we found that 2 mechanistically distinct ER stressors, thapsigargin and tunicamycin, were preferentially toxic to the MN population compared to the non-MN population, in both a dose- and time-dependent manner (Figure 1C-E). As cellular response to stress can be influenced by culture parameters, we confirmed that this preferential MN toxicity was maintained at a variety of culture densities and MN maturation statuses (Figure S1A). Moreover, exposure to the proteosome inhibitor MG132 did not result in preferential MN cell death under the conditions employed here, demonstrating that selective MN death did not result from all forms of proteostatic stress (Figure S1B-E), but rather implicated ER stress specifically in preferential MN vulnerability.

We next sought to assess whether induction of ER stress could also potentiate differential cell-type toxicity in MN cultures derived from ALS patients. We used two established ALS patient iPSC lines, each with a heterozygous familial ALS (fALS) mutation - TDP-43^G298S/+^ (47D) and SOD1^L144F/+^ (29D)(Alami et al. 2014; Boulting et al. 2011). These fALS patient iPSCs demonstrated comparable pluripotent stem cell properties and differentiation capacities to the 1016A control iPSC line (Neel et al. 2023). Furthermore, both types of fALS patient MNs also similarly displayed enhanced MN vulnerabilities to the ER stressors compared to the non-MN populations (Figure 1F). The observation that both healthy and fALS MN cultures displayed preferential MN degeneration compared to non-MN cell types, adds support to the premise that MNs are intrinsically more vulnerable to ER stressors than other cell types, a result consistent with ALS mouse and hESC stem cell models (Saxena, Cabuy, and Caroni 2009; Thams et al. 2019). Finally, both stressors could be applied to synchronously accelerate MN degeneration in healthy and ALS MN cultures (Figure 1G). While we generally observed enhanced ALS MN degeneration compared to control MN cultures (Figure 1G), line-to-line variability between iPSC derivatives excludes the conclusion that the fALS mutations themselves were sufficient to drive this phenotype. Nonetheless, simultaneously inducing MN degeneration in both healthy and fALS MN cultures enabled the development of a pharmacological human MN survival assay that encompassed genetic and nongenetic drivers of disease.

### 2.2 Early UPR signaling preceded destruction of neurites and apoptotic signaling

Thapsigargin and tunicamycin are known inducers of the UPR, a signaling pathway adapted to handle intracellular protein misfolding and hypothesized to contribute to underlying differential MN vulnerability in ALS (Kanekura et al. 2009; Matus et al. 2013; Saxena and Caroni 2011; Saxena, Cabuy, and Caroni 2009). Although UPR signaling provides an adaptive mechanism to limit ER burden and accumulation of misfolded proteins, prolonged signaling drives apoptosis via the PERK pathway.

To understand the relationship between UPR activation and initiation of preferential death in MNs, we temporally tracked UPR induction in healthy 1016A iPSC derived MNs upon exposure to thapsigargin and tunicamycin. We found that thapsigargin induced phosphorylation of eIF2α to block protein translation as soon as 15 minutes after stressor challenge, with a maximum response after 1 hour (Figure 2A). Tunicamycin addition also led to phosphorylation of eIF2α, although the activation was somewhat slower, reaching a maximum only 4 hours post-exposure (Figure 2A). We then detected splicing of XBP1 (downstream of IRE1 signaling) for thapsigargin and tunicamycin between 2 and 4 hours after treatment (Figure 2A). Upregulation of the ER resident chaperone BiP was rapidly initiated, with increases at the mRNA level between 1-4 hours after each stressor addition, and significant increases in protein levels by 8 and 24 hours (Figure 2A-B). The induction of the pro-apoptotic transcription factor CHOP displayed a similar expression pattern to BiP upregulation, with mRNA increases between 1-4 hours after each stressor treatment, and significant increases in protein levels by 8 and 24 hours (Figure 2A-B).

**Figure 2.**
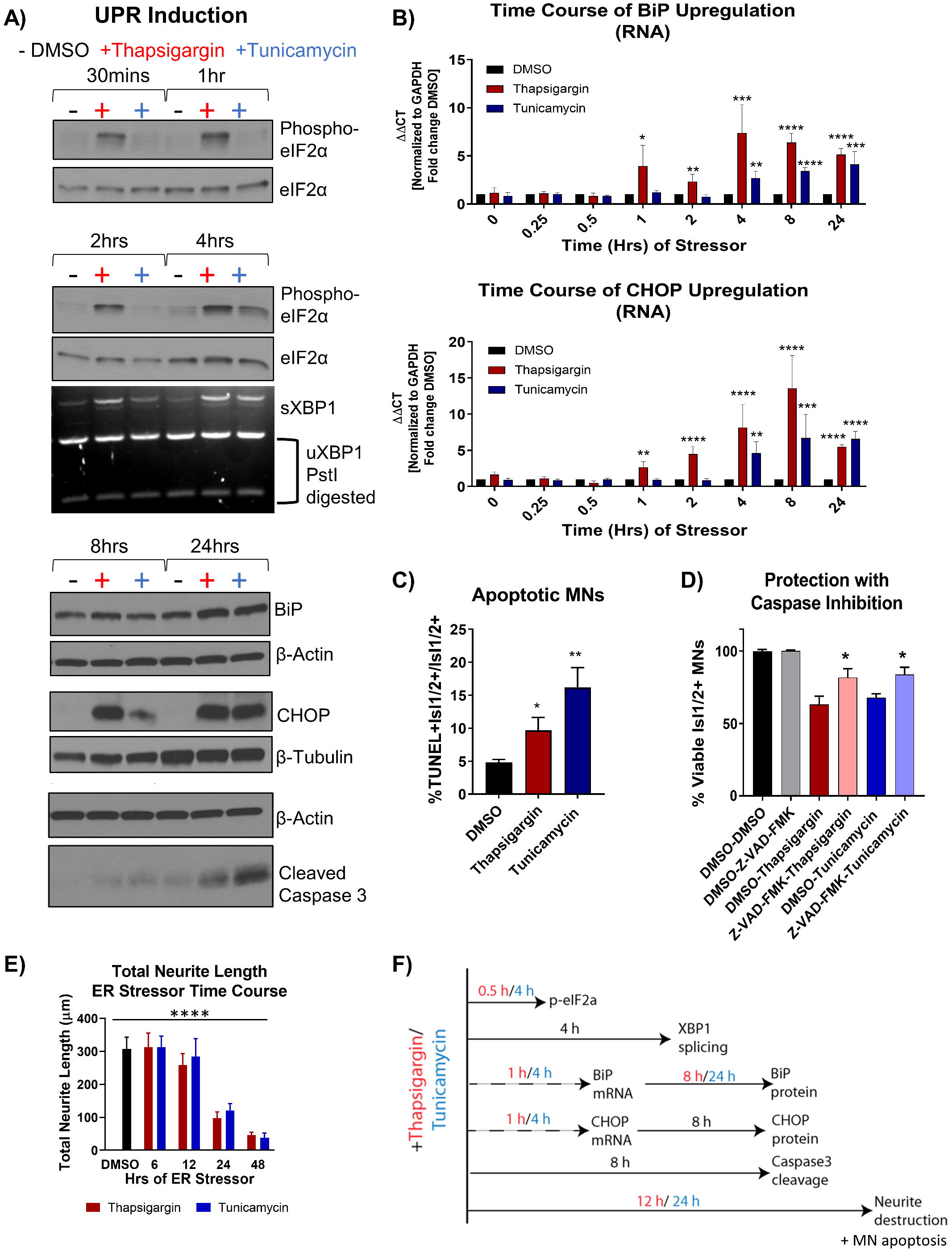
Unfolded protein response signaling drives neuritic damage and MN apoptosis. (A) Western blot and splicing-PCR analyses of patient derived MN cultures treated with either DMSO, 1μM thapsigargin or 1μM tunicamycin for various times points. Nb=3. (B) qRT-PCR analyses of patient derived MN cultures treated with DMSO, 1μM thapsigargin or 1μM tunicamycin for various times points. ΔΔCt values are graphed, normalizing target gene qPCR Ct values (BiP or CHOP) to housekeeping gene Ct values (GAPDH) and then calculating the fold change of this normalized Ct value to control conditions such that DMSO =1 at all time points. Nb=3, nt=3, p < 0.05*, <0.01**, <0.001***, <0.0001**** one-way ANOVA. (C) The percentage of apoptotic MNs in control (DMSO), 1μM thapsigargin, or 1μM tunicamycin stress conditions, quantified by the number of TUNEL+ MNs (Isl1/2+) per total MN numbers (Isl1/2+). Nb=3, nt>4, 2 tailed unpaired students t-test; Thapsigargin- p = 0.02853, Tunicamycin- p = 0.0011. (D) Quantification of MN viability in derived cultures treated with either control vehicle (DMSO), stressors alone (1μM thapsigargin or tunicamycin), or stressors with a pan-caspase inhibitor (100μM Z-VAD-FMK). Nb=3, nt >2, 2 tailed unpaired students t-test; Thapsigargin- p = 0.011583, Tunicamycin- p = 0.018997. (E) Quantification of total neurite length (μm) after exposure to 1μM ER stressors for increasing amounts of time. Nb=3, nt=6, p < 0.05*, <0.01**, <0.001***, <0.0001**** one-way ANOVA with Tukey’s hsd post-hoc test. (F) Temporal schematic of UPR-associated signaling events occurring post-treatment with thapsigargin or tunicamycin in iPSC-derived MNs. Biological replicate experiments denoted as Nb, each with technical replicate experiments nt. Data are mean value +/- SEM.

Prolonged ER stress signaling by 8-24 hour exposure to thapsigargin and tunicamycin resulted in the cleavage of caspase 3 (Figure 2A) and coincided with an increase in the percentage of TUNEL+ cells (Figure 2C). About half of the MN death caused by thapsigargin and tunicamycin treatment was also prevented by the addition of a pan-caspase apoptosis inhibitor Z-VAD-FMK, demonstrating that the selective MN toxicity was due in part to apoptosis (Figure 2D). Furthermore, apoptotic death was coincident with and often preceded by degeneration of neurites extending from the neuronal soma, which was observed with both live cell imaging (Supplemental Video Files 1-4) and high throughput neurite tracking software (Figure 2E). Taken together, these data generate a temporal guide of ER stress activation in MNs, suggesting prolonged ER stress leads to death of vulnerable MN populations via apoptosis. Additionally, these results indicate a dosing paradigm in which potential protective compounds could be evaluated for their ability to attenuate ER stress-associated toxicity. The observed temporal order of UPR-associated signaling events post-addition of thapsigargin and tunicamycin is summarized in Figure 2F.

### 2.3 Pharmacologic inhibition of MAP4K4 preserved ALS patient MN viability to a greater extent than current ALS therapeutics

We next sought to determine the extent to which the above defined ER stress MN platform could be rescued with current therapeutics. To do this, we assessed the protective effects of two FDA approved treatments for ALS, riluzole and edaravone. Interestingly, we found that none of the concentrations of riluzole and edaravone tested (10nM-10µM) protected from the toxicity caused by ER stress, as measured by the number of viable MNs and neurite architecture (Figure 3A, B). Failure of these compounds to confer MN protection could be due either to the elicited ER stress being too profound for rescue or through the distinct non-ER related mechanism of action of these drugs. To rule out that our ER stress assay was unresolvable, we performed the same experiment, but this time added kenpaullone, a kinase inhibitor found originally by our group to broadly protect murine and human MNs in various cellular culture systems (Yang et al. 2013; Reinhardt et al. 2019; Thams et al. 2019). Consistent with these previous studies, we found that treatment with kenpaullone dramatically improved human MN survival in response to both ER stressors, in healthy control 1016A, TDP-43, and SOD1 fALS patient iPSC-derived MNs (Figure 3C). This preservation of MN viability coincided with the maintenance of TUJ1+ neuritic networks (Figure 3D). As MN death via thapsigargin and tunicamycin could be attenuated, these data imply that riluzole and edaravone were ineffective in these assays, consistent with recent work investigating the effectiveness of riluzole and edaravone in rescuing ALS toxicity in similar patient-derived cell culture models (Fujimori et al. 2018; Yang et al. 2013).

**Figure 3.**
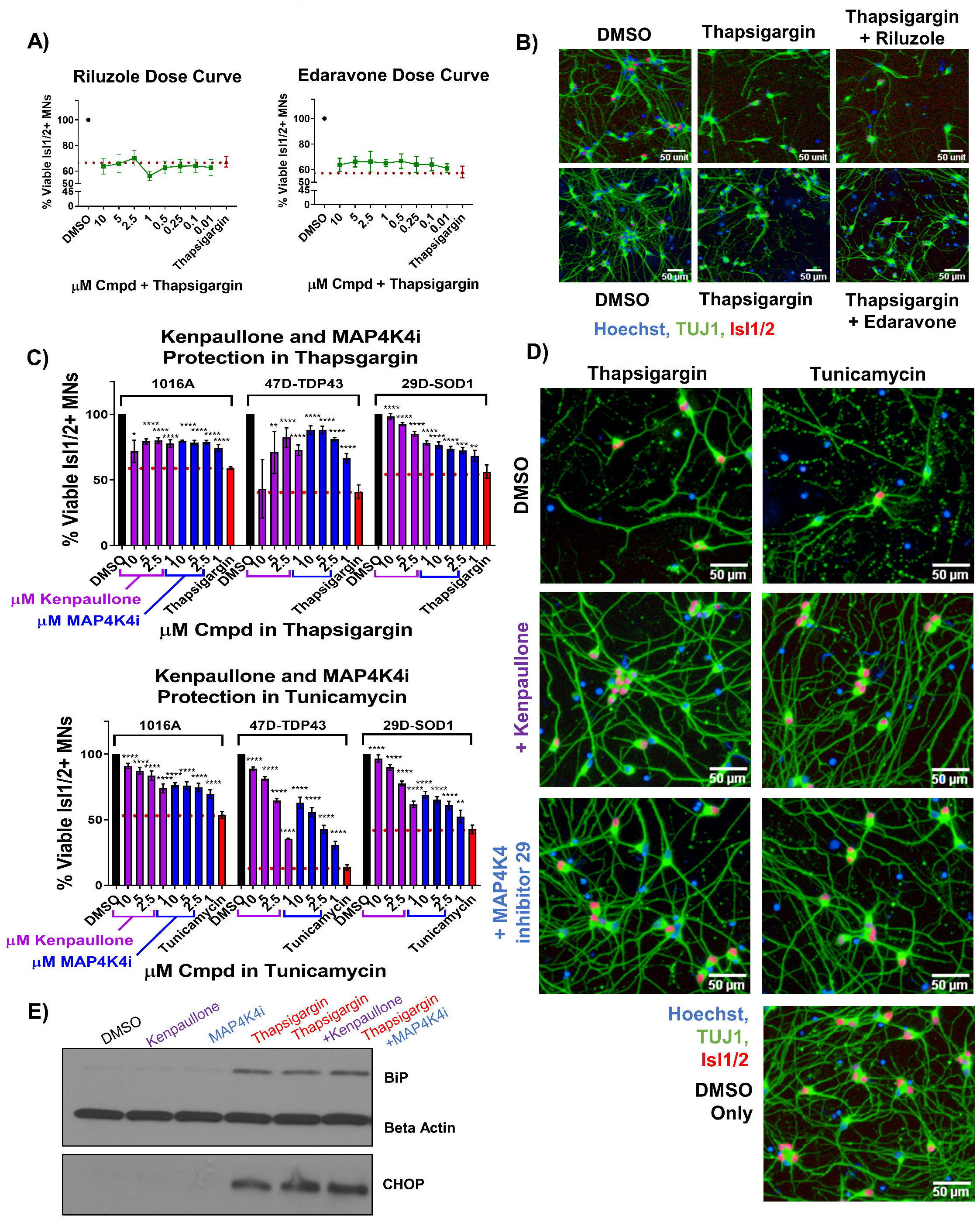
MAP4K4 inhibitors preserve MN viability better than the current FDA approved ALS drugs without reducing UPR induction. (A) Quantification of MN viability in patient derived cultures treated for 48hrs with various doses of Riluzole or Edaravone in 1μM thapsigargin. Nb=3, nt=3. (B) Representative immunofluorescent staining of patient derived MNs in control (DMSO), thapsigargin, or Riluzole/Edaravone thapsigargin treated conditions. (C) Quantification of healthy patient MN viability (1016A) or ALS patient MN viability (47-TDP-43^G298S^ mutant or 29D-SOD1^L144F^ mutant) after 48hrs treatment with various doses of kenpaullone or MAP4K4 inhibitor 29 in ER stress conditions (1μM thapsigargin or tunicamycin). Nb=3, nt=3, p < 0.05*, <0.01**, <0.001***, <0.0001**** one-way ANOVA with Tukey’s hsd post-hoc test. (D) Representative immunofluorescent images of ALS TDP-43^G298S^ mutant patient derived MN cultures in control (DMSO), ER stress, or ER stress and kenpaullone/MAP4K4 inhibitor 29 rescued conditions. (E) Western blot analyses of BiP and CHOP in patient derived MN cultures treated with either DMSO vehicle, 3μM kenpaullone, 5μM MAP4K4i, 1μM thapsigargin, or kenpaullone/MAP4K4i in thapsigargin. Nb=3. Biological replicate experiments denoted as Nb, each with technical replicate experiments nt. Data are mean value +/- SEM.

As kenpaullone has multiple cellular targets (Leost et al. 2000; Yang et al. 2013) and is unlikely to be CNS penetrant (Kitabayashi et al. 2019), we aimed to gain further insight into its protective effects in the hopes of identifying additional, druggable targets for MN degeneration in ALS. Previous work indicated that MAP4K4 was among the kinases inhibited by kenpaullone (Yang et al. 2013) and that inhibition of MAP4K4 alone was sufficient to protect against trophic factor withdrawal-induced ALS MN degeneration (Wu, Watts, and Rubin 2019). Therefore, we next tested whether treatment with a more selective compound inhibitor of MAP4K4 (MAP4K4 inhibitor 29, (Crawford et al. 2014) would also confer protection from ER stress. We found that MAP4K4 inhibitor 29 indeed protected viability and neurite morphology in response to both ER stressors, in control 1016A, TDP-43, and SOD1 fALS patient MNs, similar to kenpaullone (Figure 3C, D). In addition to confirming the robustness of our platform for modeling ER stress-associated MN toxicity and finding neuroprotective drugs, these results provide further evidence that inhibition of MAPK4K is protective of MNs.

### 2.4 Pharmacological MAP4K4 inhibition protected MNs against ER stress-associated toxicity despite UPR induction

Since exposure to thapsigargin and tunicamycin rapidly upregulates UPR-associated signaling pathways eventually leading to cell death, we sought to explore whether kenpaullone and MAP4K4 inhibition protected against ER stress by directly blocking these signaling cascades. To test this, we exposed iPSC-derived MNs to thapsigargin in the presence or absence of kenpaullone or MAP4K4 inhibitor 29 and evaluated protein level expression of chaperone BiP and stress-induced transcription factor CHOP via Western blot. We observed sustained upregulation of both BiP and CHOP in cultures treated with kenpaullone or the MAP4K4 inhibitor (Figure 3E), consistent with a mechanism independent of blocked UPR induction.

### 2.5 Phosphoproteomic analyses identified JNK, PKC and BRAF as convergent cellular perturbations of protective MAP4K4 inhibitors

Results thus far demonstrated the ability of kenpaullone and a MAP4K4 inhibitor to protect against MN death under various conditions, corroborating Wu et al. 2019 (Wu, Watts, and Rubin 2019). However, MAP4K4 shares a high degree of structural similarity to other upstream MAP kinases, including MINK1, with nearly 100% sequence homology at the kinase ATP-binding pocket. Moreover, studies have demonstrated that small molecule inhibitors of MAP4K4 also target additional kinases including MINK1 and TNIK (Bos et al. 2019; Larhammar, M., Huntwork-Rodriguez, S., Rudhard, Y., Sengupta-Ghosh, A. & Lewcock 2017). As a result, we sought to further define the landscape underlying pharmacological MN protection conferred by MAP4K4 inhibition in the presence of prolonged ER stress. To do so, we performed global phosphoproteomic analyses on patient MN cultures treated with thapsigargin in the presence or absence of kenpaullone or MAP4K4 inhibitor 29 (Figure S2A). A moderate dose of thapsigargin was used, and total protein lysates from MN cultures were collected at early and late time points (24 and 48 hours, respectively) (Figure S2A), allowing a dynamic view of cellular perturbations occurring with time while ensuring relevant molecular changes were captured in surviving cells (Figure S2A).

From 2 biological replicate experiments, a total of 67,357 and 59,708 peptides, and 28,399 and 28,973 phosphopeptides were quantified. This corresponded to 6,697 unique proteins, 2,764 unique phosphoproteins and 7,999 nonredundant phosphorylation sites, with the changes in phospho-site abundance being normalized to protein levels when possible (Table S1, Supplementary Data Set 1). We first observed that thapsigargin treatment alone drove the largest alterations in the proteome and phosphoproteome (Figure 4A, S2B). Significantly upregulated proteins in thapsigargin treated conditions included the ER chaperone BiP (HSPA5) and the pro-apoptotic protein Bcl-2 binding protein 3 (BBC3), consistent with our results showing thapsigargin induces UPR-driven apoptosis in human MNs (Figures 2, 4A). Gene ontology analysis of significantly upregulated proteins (FDR<0.05) in thapsigargin-stressed cells revealed enriched terms including “response to endoplasmic reticulum stress” and “positive regulation of cytosolic calcium ion concentration” (Figure 4B), consistent with known effects of thapsigargin as an inhibitor of the ER calcium ATPase pump (cytosol to ER calcium transporter).

**Figure 4.**
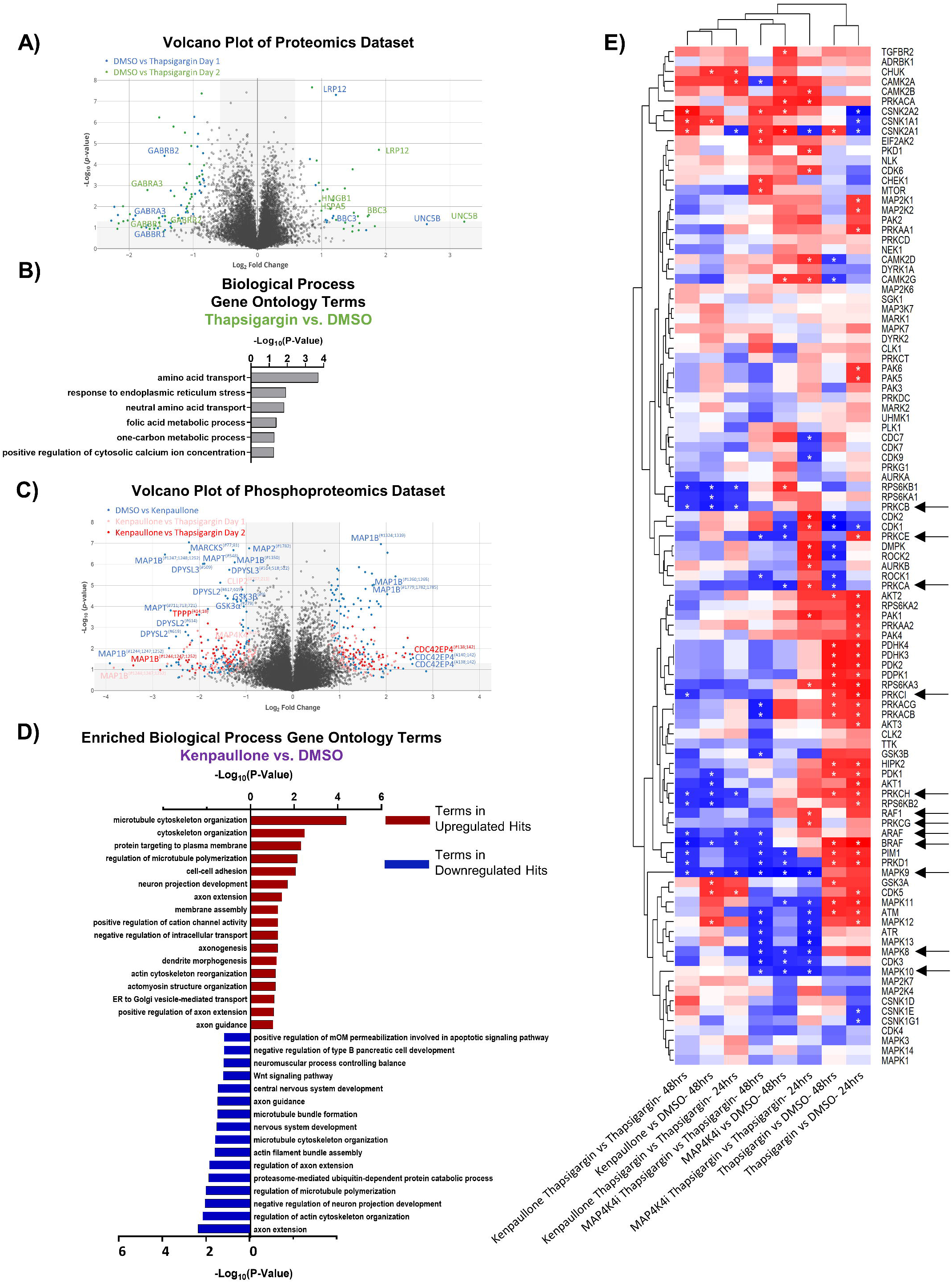
Proteomic and phosphoproteomic analyses of degenerating MNs protected with MAP4K4 inhibitors. (A) Volcano plot of all proteins quantified in the proteomics dataset. Individual proteins are represented as single dots. (B) Gene ontology analyses of significantly upregulated proteins in response to thapsigargin treatment. (C) Volcano plot of all phosphoproteins quantified in the phosphoproteomics dataset. Individual phosphoproteins are represented as single dots, with round shapes indicating normalized phosphoprotein levels (to protein levels), and diamond shapes indicating non-normalized phosphoprotein levels (due to lack of non-phosphorylated protein quantification). Selected phosphoproteins are annotated with the phosphorylation site position indicated as a superscript. (D) Gene ontology analyses of the significantly upregulated phosphoproteins or downregulated phosphoproteins in response to kenpaullone treatment. (E) Unbiased hierarchical clustering of kinases whose phosphosubstrates were perturbed with compound treatments (identified by kinase-substrate enrichment analyses (KSEA)). Red boxes indicate upregulated kinase substrates; blue boxes indicate downregulated kinase substrates. Stars indicate FDR q values < 0.05.

In addition to thapsigargin, we observed that kenpaullone exposure significantly perturbed the phosphoproteome (Figure 4C, S2B). As expected, known targets of kenpaullone, including phosphorylated MAP4K4 and GSK3β/α, were notably decreased with kenpaullone treatment alone (Figure 4C). Moreover, gene ontology terms of the significantly dysregulated proteins from kenpaullone treatment largely indicated changes in neuron extension processes, including: “positive regulation of axon extension”, “neuron projection development”, “dendrite morphogenesis’, and “microtubule cytoskeleton organization” (Figure 4D), with the main drivers of these changes including the microtubule plus-end tracking protein CLASP2, the neuronal guidance microtubule assembly protein DPYSL2, and the microtubule associated proteins MAP1B and MAPT (Figure 4C). Since cultures treated with the more selective MAP4K4 inhibitor 29 did not exhibit changes in these microtubule proteins, it is likely that these phosphoproteomic changes were due to kenpaullone’s dual inhibition of GSK3β and CDK5, rather than MAP4K4 inhibition, consistent with previously published work (Reinhardt et al. 2019). Downregulation of “apoptotic signaling pathways” was also observed in the gene ontology terms (Figure 4D), further supporting the pro-survival effect observed with kenpaullone treatment.

We next utilized pathway enrichment analyses to interrogate the mechanisms underlying the protection from thapsigargin exposure conferred by MAP4K4 inhibitor 29. Unbiased hierarchical clustering of kinase-substrate enrichment analyses revealed several kinase pathways commonly downregulated by both kenpaullone and MAP4K4 inhibitor 29 (Figure 4E), including an expected decrease in the JNK pathway (annotated as MAPK8/9/10), a known downstream pro-apoptotic target of MAP4K4 signaling. Additionally, several unexpected targets were commonly altered between kenpaullone and MAP4K4 inhibitor 29 treatment, including protein kinase family members (PRKD1, PRKCG, PRKC1, PRKCB, PRKCE) as well as RAFs (BRAF, ARAF, RAF1), suggesting that these proteins might also play a neuroprotective role in MNs.

### 2.6 Small molecule inhibitors to JNK, PKC, and BRAF prevented ER stress-associated MN death

To assess if the shared pathways dysregulated by kenpaullone and MAP4K4 inhibitor 29 were functionally consequential in MN survival, we tested selective small molecule inhibitors of JNK (SP600125), PKC (Enzastaurin), and BRAF (GDC-0879) in an 8-point dose response survival curve on patient MNs treated with thapsigargin stress. In line with our work and that of others implicating JNK signaling in ALS MN degeneration (Wu, Watts, and Rubin 2019; Bhinge et al. 2017), we observed that the JNK inhibitor SP600125, as well as the PKC inhibitor Enzastaurin and the BRAF inhibitor GDC-0879, all preserved MN viability and TUJ1+ neurite networks to levels comparable to those obtained with kenpaullone or MAP4K4 inhibitor 29 (Figure 5A-B). These small molecule inhibitors also significantly protected MN numbers and neurite morphology in the presence of tunicamycin stress (Figure 5C-D), indicating the protection was not thapsigargin-specific.

**Figure 5.**
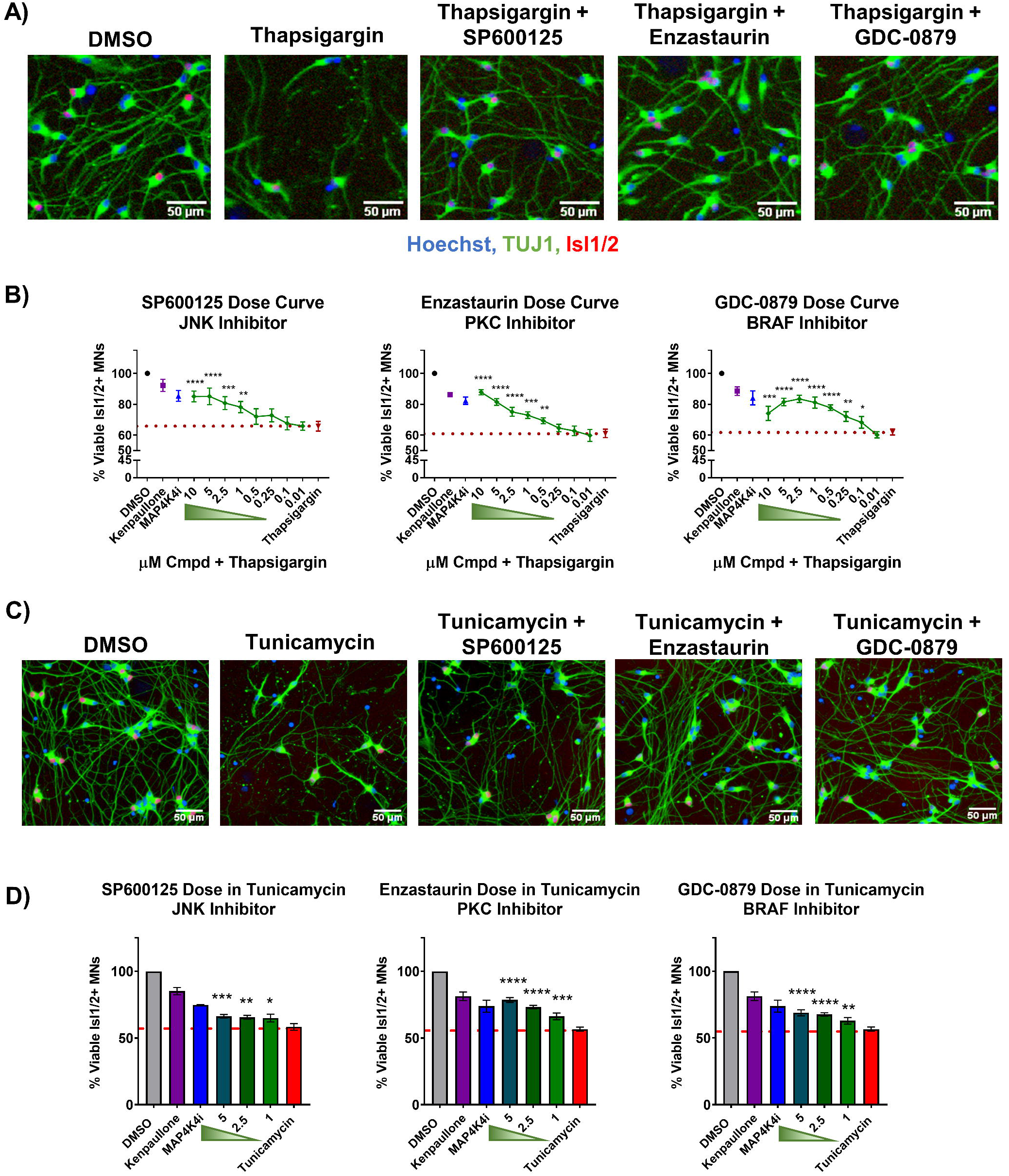
Small molecule inhibitors to JNK, PKC, and BRAF are MN protective compounds. (A) Representative immunofluorescent images of patient derived MN cultures in control (DMSO), thapsigargin stress, or small-molecule-inhibitor-rescued conditions. (B) Quantification of MN viability in patient derived cultures after 48hrs treatment with various doses of hit small molecule inhibitors in 1μM thapsigargin. 3μM kenpaullone or 5μM MAP4K4i in thapsigargin stress were used as positive controls, DMSO vehicle as a negative control. Nb≥3, nt=3, p < 0.05*, <0.01**, <0.001***, <0.0001**** one-way ANOVA with Tukey’s hsd post-hoc test. (C) Representative immunofluorescent images of patient derived MN cultures treated with control vehicle (DMSO), 1μM tunicamycin, or 1μM tunicamycin with 5μM small molecule-inhibitors. (D) Quantification of MN viability in patient derived cultures after 48hrs treatment with various doses of hit small molecule inhibitors in 1μM tunicamycin. 3μM kenpaullone or 5μM MAP4K4i in tunicamycin stress were used as positive controls, DMSO vehicle as a negative control. Please note that values for tunicamycin, kenpaullone and MAP4K4i rescue in Enzastaurin and GDC-0879 dose curves are the same as these experiments were performed on the same plate. Nb=3, nt=3, p < 0.05*, <0.01**, <0.001***, <0.0001**** one-way ANOVA with Tukey’s hsd post-hoc test. Biological replicate experiments denoted as Nb, each with technical replicate experiments nt. Data are mean value +/- SEM.

## 3 Discussion

ALS is an extremely complex MN disease with few therapeutic options. Recent advances in generating MNs from healthy control and ALS patient iPSCs have greatly helped address limitations of traditional rodent models and improved drug discovery by enabling scalable production of affected cells for high-throughput drug screening (Dimos et al. 2008; Fujimori et al. 2018). Here, we sought to build upon traditional iPSC models of ALS by employing two pharmacological models of ER stress in ALS patient MN cultures, and further leveraged this platform for large-scale proteomics analysis and high-content neuroprotective drug discovery. We specifically demonstrated that the ER stressors synchronously recapitulate early disease signaling and late-stage ALS associated degenerative phenotypes of preferential MN death, in both healthy control and ALS patient MN cultures. We further found that these culture systems could predict neuroprotective targets discovered in other ALS models, and identified several novel neuroprotective agents, including a PKC inhibitor Enzastaurin and an FDA approved BRAF inhibitor GDC-0879.

One hallmark of ALS sometimes overlooked in cell culture models, includes differential MN vulnerability, where upper MNs in the motor cortex of the brain and lower MNs in the brainstem and spinal cord are disproportionally damaged in patients relative to other CNS cell types (Ferraiuolo et al. 2011; Ragagnin et al. 2019). To recapitulate this phenomenon in our model, we employed a MN differentiation protocol known to produce MNs as well as non-MN cell types of LMC identity, including interneurons and glia. Consistent with our reasoning, utilizing the heterogeneous culture of cells produced by this differentiation protocol, we observed that 2 mechanistically distinct ER stressors were both preferentially toxic to MNs compared to non-MN cell types in healthy and ALS patient cultures, demonstrating our model recapitulates this differential sensitivity and supports the proposition that perturbed ER homeostasis underlies intrinsic MN vulnerability in disease. Furthermore, as this MN selective phenotype was not observed with MG132, this indicates that different forms of proteostatic stress result in differential MN sensitivities.

As ALS is commonly sporadic with complex environmental and age-related dysfunctions significantly contributing to disease, it was important that selective vulnerability of MNs not only be observed in ALS MNs but in healthy control MNs as well. We reasoned that inducing human MN degeneration, even in MNs not affected with a defined fALS mutation, would aid in the identification of neuroprotective agents that would be effective against multiple forms of MN disease (Yang et al. 2013). Demonstrating that thapsigargin and tunicamycin universally caused an ALS-associated phenotype of preferential MN degeneration in all human MN cultures in an accelerated and reproducible timeline enabled development of a drug discovery platform compatible for high throughput drug screening assays compared to extended culture systems typically used.

To be useful for ALS drug discovery, screening-based assays must demonstrate an ability to predict neuroprotective drugs. We demonstrated this potential in our ER stressor assays using kenpaullone, a published MN protective agent identified by our group (Yang et al. 2013) and validated by others (Reinhardt et al. 2019; Thams et al. 2019), as well as a small molecule inhibitor to MAP4K4, one of kenpaullone’s targets (Crawford et al. 2014; Yang et al. 2013). We found that both compounds were able to increase control, SOD1 and TDP-43 fALS MN viability and neurite morphology, and that this rescue was greater than that of riluzole or edaravone, 2 FDA approved drugs for ALS. The inefficacy of riluzole and edaravone to confer neuroprotection is important to note. Our results are indeed largely consistent with those from the Okano group (Fujimori et al. 2018; Yang et al. 2013) who also demonstrated that these ALS treatments failed to protect against ALS MN neurite regression, cytotoxicity, and FUS+ or phosphorylated-TDP-43+ protein aggregates following extended culture maturation times (∼ 40-70 days). The inefficacy of both compounds in these platforms is unlikely due to a limitation of either assay to detect protective compounds with mechanistically distinct actions. Since riluzole and edaravone both only modestly prolong patient survival and improve motor function, these data leave open the possibility that drugs with more robust in vitro MN protective activity may benefit patients more than the existing ALS treatments (Andrews et al. 2020; Samadhiya et al. 2022; Witzel et al. 2022).

In this study, we combined the scalability of human stem cell-based MN culture with mass spectrometry to capture the entire proteomic and phosphoproteomic temporal landscape in MNs exposed to degenerative stress or protective small molecules. In doing so, we observed expected molecular changes, including decreased JNK and cJUN signaling with both MN protective agents kenpaullone and MAP4K4 inhibitor 29, and that pharmacological reduction of this signaling pathway with a JNK inhibitor (SP600125) was protective. Moreover, we discovered that protein kinase family members (PRKD1, PRKCG, PRKC1, PRKCB, PRKCE) and RAFs were also significantly dysregulated with kenpaullone and MAP4K4 inhibitor 29, and identified Enzastaurin (a PKC inhibitor), and GDC-0879 (a B-RAF inhibitor), as protective in our ER stress MN assay. Enzastaurin has been determined to be well-tolerated in clinical trials (Bourhill, Narendran, and Johnston 2017; Shuo and Steven T 2007), and GDC-0879 is already in clinical development for certain cancers, highlighting the notion that current drugs may be re-purposed for ALS therapeutics.

In a variety of studies, PKC and BRAF have been associated with neuronal viability in a context-dependent manner (Frebel and Wiese 2006; Kolch 2001; Nishizuka 1986; Tanaka and Koike 2001; Wiese et al. 2001; Zhu et al. 2004). Here, in ALS patient-specific iPSC-derived MNs, we observed a neuroprotective effect for PKC inhibition via Enzastaurin, that we speculate to be via activation of the AKT pro-survival pathway based on data from rat cerebellar granule cell neurons (Zhu et al. 2004). BRAF is a proto-oncogene with reported roles in embryonic MN and sensory neuron survival in response to neurotrophic factors (Davies et al. 2002; Wiese et al. 2001). Compound inhibitors to BRAF, including GDC-087, target the BRAFV600E/K tumors, exhibiting paradoxical activation of the MEK/ERK pro-survival pathway in cells with BRAFWT via RAF transactivation (Agianian and Gavathiotis 2018; Sieber et al. 2018; Uenaka et al. 2018). Here, we speculate that BRAF inhibition with GDC-0879 protects patient MNs challenged with ER stress due to restored activation of MAPK/ERK pro-survival signaling.

Ultimately, these data demonstrate that pharmacologic ER stress induces MN specific toxicity in iPSC-derived cultures by initiating the UPR and apoptosis (Figure 6). By combining this model of MN toxicity with proteomics and phosphoproteomics, we identified druggable targets capable of protecting ALS MNs from ER stress-associated toxicity, including PKC and BRAF signaling components (Figure 6). Collectively, this work outlines a platform to exacerbate ER stress in patient derived MNs in a scalable manner to identify novel neuroprotective compounds.

**Figure 6.**
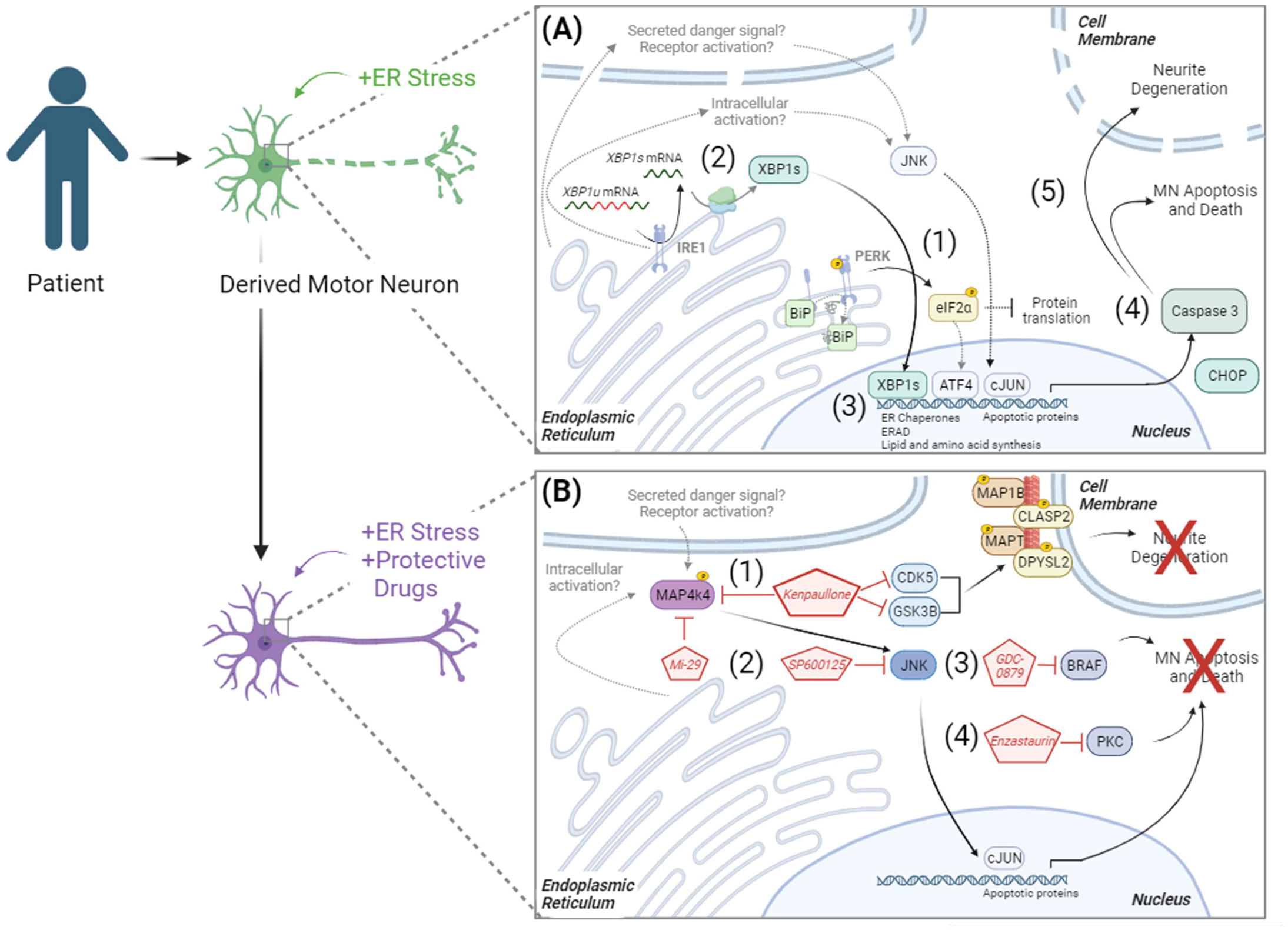
Model of ER stress induced preferential MN degeneration and pharmacological protection. ER stress induced preferential MN degeneration in both healthy and ALS patient MN cultures, and small molecule inhibitors to intracellular signaling pathways could protect against this MN degeneration. (A) In conditions of ER stress, the unfolded protein response was activated, with (1) PERK-mediated phosphorylation of eIF2a and (2) IRE1-mediated splicing of XBP1 upregulating protein folding chaperones BIP (3), ER associated degradation (ERAD) proteins, and lipid and amino acid biosynthesis. With chronic/prolonged UPR signaling, apoptotic proteins like CHOP were upregulated and caspase 3 was cleaved (4), driving neurite degeneration and MN apoptosis (5). (B) In the presence of neuroprotective compounds, ER stress induced MN death was attenuated. (1) Kenpaullone mediated MN protection through the dual inhibition of GSK3β and CDKs, which reduced the aberrant hyperphosphorylation of microtubule associated proteins, as well as through the inhibition of MAP4K4. (2) MAP4K4 inhibition by kenpaullone or the more selective MAP4K4 inhibitor 29 (Mi-29) promoted MN protection by attenuating the pro-apoptotic JNK/cJUN pathway. Specific inhibition of JNK with SP600125 additionally conferred MN protection. PKC and BRAF were additional proteins commonly dysregulated by the protective agents kenpaullone and MAP4K4 inhibitor 29, and a BRAF inhibitor GDC-0879 (3) and a PKC inhibitor Enzastaurin (4) also functionally promoted MN protection. The neuroprotective mechanism underlying Enzastaurin and GDC-0879 remains to be experimentally determined. Moreover, how MAP4K4 is activated in ER stress and MN death conditions such that its inhibition is neuroprotective also remains to be determined. We hypothesize either an intracellular activation mechanism and/or an extracellular activation mechanism via a secreted protein or receptor activation. Proteins and pathways experimentally determined in this publication are colored with black arrows. Proteins and pathways not experimentally detected in this publication, but with literature support, are non-colored with dashed grey arrows.

## 4 Materials and Methods

### 4.1 hiPSC culture

All human induced pluripotent stem cell (hiPSC) culture was performed with approval by the institutional review board and the Harvard Committee on the Use of Human Subjects. hiPSC cultures were maintained at 37°C with 5% CO_2_ in a ThermoFisher Scientific biological incubator. hiPSCs were cultured with supplemented StemFlex medium (ThermoFisher Scientific) on Matrigel-coated (BD Biosciences) tissue culture plates. Cells were fed every other day and passaged at ∼80% confluency using 5 minutes of room temperature 0.5mM EDTA (Life Technologies) followed by mechanical disruption with a cell lifter. All cell lines were confirmed genotypically correct, karyotypically normal by analysis of 20 metaphase spreads by Cell Line Genetics or WiCell Cytogenetics, and free of mycoplasma contamination using the LookOut Mycoplasma PCR Detection Kit (Sigma Aldrich) or the MycoAlert PLUS Mycoplasma Detection Kit (Lonza).

### 4.2 MN differentiation and dissociation

hiPSCs were differentiated into MNs as described previously (Maury et al. 2014; Neel et al. 2023). Briefly, confluent (60-90%) hiPSC cultures were first detached from Matrigel (BD Biosciences) coated plates and dissociated into single cells using 37°C Accutase (Stem Cell Technologies) for ∼5 minutes. Accutase was quenched and single cells were seeded for differentiation in suspension at a density of 1×10^6^ cells/mL in complete mTeSR media (Stem Cell Technologies) supplemented with 10ng/mL FGF2 (Peprotech) and 10µM ROCK-inhibitor Y-27632 (Stemgent). 24 hours after seeding, cells were filtered through a 100µm cell strainer (VWR) and additional, equal volume complete mTeSR media (Stem Cell Technologies) was added. 48 hours after seeding (differentiation day 0), mTeSR media was replaced with N2B27 MN differentiation media, composed of a v:v mixture of DMEM/F12 and Neurobasal media (Life Technologies), supplemented with 1% N2 (Life Technologies), 2% B27 (Life Technologies), 1% Pen-Strep (Life Technologies), 1% Glutamax (Life Technologies), 0.1% beta-mercaptoethanol (βME, Life Technologies), and 20µM ascorbic acid (Sigma Aldrich). Day 0 and day 1 of differentiation, N2B27 MN differentiation media was supplemented with 10µM SB-431542 (R&D Systems), 100nM LDN-193189 (ReproCELL), and 3µM CHIR-99021 (ReproCELL). Day 2 and Day 4, differentiation media was supplemented with 10µM SB-431542, 100nM LDN-193189, 3µM CHIR-99021, 1µM retinoic acid (Sigma Aldrich), and 1µM smoothened agonist (DNSK International, LLC). Day 5, differentiation media was supplemented with 1µM retinoic acid, and 1µM smoothened agonist, Day 7 with 1µM retinoic acid, 1µM smoothened agonist, and 20ng/mL brain derived neurotrophic factor (BDNF, R&D Systems). Day 9, differentiation media was supplemented with 1µM retinoic acid, 1µM smoothened agonist, 20ng/mL BDNF, and 10µM DAPT (R&D Systems). Day 11 and 13, differentiation media was supplemented with 1µM retinoic acid, 1µM smoothened agonist, 20ng/mL BDNF, 10µM DAPT, and 20ng/mL glial derived neurotrophic factor (GDNF, R&D Systems).

Day 15 of differentiation, embryoid bodies (EBs) were collected, washed once with 1x phosphate buffer solution (PBS) without calcium and magnesium (VWR), and dissociated with 0.25% Trypsin-EDTA (Life Technologies) and 50µg/mL DNase1 (Worthington Biochemical) for 5 minutes at 37°C with movement. Trypsin was quenched with fetal bovine serum (FBS, Sigma Aldrich), and centrifuged at 400g for 5 minutes. The cell pellet was then resuspended in dissociation buffer, consisting of 5% fetal bovine serum (FBS, Sigma Aldrich), 25mM glucose, 1% Glutamax, in 1x PBS without calcium and magnesium, and mechanically triturated using a p1000 pipet. Dissociated single cells were pelleted by centrifugation (400g, 5 minutes) and resuspended in complete MN media, consisting of Neurobasal media supplemented with 1% N2, 2% B27, 1% Pen-Strep, 1% Glutamax, 1% Non-essential amino acids (Life Technologies), 0.1% βME, 20µM ascorbic acid, 20ng/mL BDNF, GDNF, and CNTF (ciliary neurotrophic factor, R&D Systems), and 10µM UFDU (v:v Uridine (Sigma Aldrich):Fluorodeoxyuridine (Sigma Aldrich)). Resuspended dissociated single cells were filtered through a 40µm cell strainer, counted with a 1:1 trypan blue dilution using an automated cell counter, and plated at the desired density on tissue culture treated plates coated with 1X borate buffer (Life Technologies), 25µg/mL poly-ornithine (Sigma Aldrich), 5µg/mL mouse laminin (Life Technologies), and 10µg/mL fibronectin (VWR).

### 4.3 ER stressor assays

For survival analyses, dissociated MNs were plated in complete MN media at a density of 50,000 cells/well (unless otherwise indicated) in the inner 60 wells of borate/poly-ornithine/laminin/fibronectin coated 96-well plates (Perkin Elmer). Outer wells were filled with water to avoid evaporation effects. 3 days after plating, ¾ media was removed and replaced with fresh complete MN media. 6 days after plating, all media was removed, and MN cultures were treated simultaneously with ER stressor media or ER stressors with protective compounds. Unless otherwise indicated in dose response curves, 1µM of thapsigargin (Sigma Aldrich) or 1µM tunicamycin (Sigma Aldrich) were used for standard stressor conditions, and 3µM of kenpaullone (Tocris) and 5µM of MAP4K4 inhibitor 29 (Genentech) in 1µM ER stressor media were protective positive controls. 0.1% DMSO (Sigma Aldrich) complete MN media was the negative control, and total concentration of DMSO was equal to 0.1% in all wells. MNs were incubated with stressors, protective compounds, and test compounds, for 48 hours and fixed with 4% paraformaldehyde (PFA, VWR) prior to quantitative analysis of Isl1/2 (Abcam), TUJ1 (Biolegend) and Hoechst (Life Technologies) staining as described in Immunofluorescent staining and high content image analysis methods section. All cell counts were expressed as a percentage of surviving DMSO-control cells.

For gene expression analyses, dissociated MNs were plated in complete MN media at a density of 2×10^6^ cells/well of 6-well plates coated with borate, poly-ornithine, laminin, and fibronectin. 3 or 6 days after plating, media was removed and MN cultures were treated with 1μM thapsigargin, tunicamycin, MG132 stressor media, alone or with 3µM of kenpaullone (Tocris) or 5µM of MAP4K4 inhibitor 29 (Genentech). Equal concentration DMSO was used as a negative control. Samples were collected at the appropriate timepoints for RNA and protein expression analyses as described in Protein extraction and western blotting and RNA isolation, reverse transcription, and quantitative PCR methods sections.

### 4.4 Immunofluorescent staining and high content image analysis

Cells were fixed for 15 minutes at room temperature with 4% paraformaldehyde (PFA, Sigma Aldrich) in PBS, achieved by adding equal volume 8% PFA to equal volume culture media. Fixed cells were gently washed once with PBS, then blocked with 10% normal goat serum, 0.1% Triton X-100 in 1x PBS for 30 minutes at room temperature. Blocked cells were then incubated for 1 hour at room temperature with primary antibodies anti-Isl1/2 (Abcam ab109517, 1:2000), anti-Chx10 (Santa Cruz Biotechnology sc-365519, 1:50), and anti-TUJ1 (Biolegend 801202, 1:2000, or Novus Biologicals NB100-1612, 1:2000). Following primary antibody incubation, cells were gently washed once with PBS, and incubated for 1 hour at room temperature with 2µg/mL Hoechst (Life Technologies H3569) and species-matched, fluorophore-conjugated secondary antibodies (Life Technologies Alexa-488, -546, -555, or -647, 1:1000) diluted in 10% normal goat serum, 0.1% Triton X-100, in PBS. Immunofluorescent labeled cells were gently washed twice with PBS before image acquisition.

High content screening systems (Operetta (PerkinElmer) or ImageXpress (Molecular Devices)) were used for all image acquisition. Images were acquired automatically using a 10x or 20x objective, a solid-state laser light source, and a sCMOS (scientific complementary metal-oxide-semiconductor) camera, corresponding to 9-12 evenly distributed fields per well of each 96-well plate. Using these parameters, at least 2,000 cells were imaged per well of each 96-well plate, with typically >5,000 cells/well with normal assay conditions. Quantitative analyses of these images were then performed using the Columbus/Harmony image analysis software (PerkinElmer). Manually designed scripts were written to define a viable cell population, which consisted of intact, Hoechst-stained nuclei larger than ∼45µm^2^ (range 37-55) in surface area and with intensities lower than the threshold brightness of pyknotic nuclei (Figure S3A-B). From this viable cell population, the neuron-specific βIII-Tubulin marker was used to define the neuronal population and neurite morphologies (as described below, Figure S4). From the viable population, nuclear Isl1/2 antibody staining was used to define the MN population, using intensity thresholds that accurately reflect positive nuclear staining (Figure S3C). With these set parameters, total numbers of nuclei, viable neurons, and viable MNs were then quantified automatically across the plate, ensuring unbiased measurements for all test conditions. Images were visually inspected during analysis and script generation, and after analysis to ensure data validity. All cell counts were then expressed as a percentage of surviving DMSO-control cells.

For neurite detection, the CSIRO neurite analysis 2 method (Harmony/Columbus, PerkinElmer) was used on βIII-Tubulin (TUJ1) staining of total Hoechst+ nuclei populations. This generated a mask that traced and segmented neuritic processes extending from individual neurons (Figure S4). Scripts were optimized per plate to achieve accurate neurite tracking, with the following parameters typically used-smoothing width at ∼2-3px was used to suppress noise and obtain a single maximum intensity across the neurites with gaussian filtering; linear windows at ∼11px specified the dimension in pixels used to find local βIII-Tubulin intensity maxima and contrast parameters between 1-2.5 were used to decrease background noise. Small, spurious objects were eliminated by removal of small diameter objects >3px, and extraneous lateral projections were cleaned using a debarb length of <9-15px. Gap closure distance and tree length were optimized to ensure gaps between detected neurites were closed, without making false connections, and that neurite bodies were linked to their corresponding mother cell. All output counts were the values per neuron, averaged per well.

Representative images displayed in figures were cropped using FIJI/ImageJ. All automatic contrast settings in Harmony/Columbus were first disabled. Images were then selected for each condition, saved from the database, and imported into FIJI/ImageJ. A region of interest (ROI) was generated and used for each image per condition, allowing the preservation of equal image scale. Cropped images were then saved as final high-resolution TIFs.

### 4.5 Automated live cell imaging

Dissociated MNs were plated in complete MN media at a density of 50K, 25K, and 12.5K cells/well in the inner 60 wells of borate/poly-ornithine/laminin/fibronectin coated 96-well plates (Greiner). Outer wells were filled with water to avoid evaporation effects. 3 days after plating, ¾ of the media was removed and replaced with fresh complete MN media. 6 days after plating, and before treatment, the MN culture plate was entered into the Nikon BioStation CT for an initial image acquisition. A 10x objective was used to acquire phase images across the plate with a 4 x 4 stitched tiling capture area equivalent to 3.08 x 3.08mm per well. After an initial image acquisition, the plate was removed, and MN culture media was replaced with stressor media as indicated in ER stressor assays methods section. The plate was then returned to the Nikon BioStation CT for image acquisition every 6 hours for 48 hours. Final images were then saved as time-lapse video files using CL-Quant software (Nikon) and FIJI/ImageJ was used to select and crop a region of interest for representative video files.

### 4.6 Protein extraction and western blotting

Protein from 2×10^6^ cells was harvested on ice after 1 PBS wash using the Pierce RIPA lysis and extraction buffer (25 mM Tris-HCl, pH 7.6, 150 mM NaCl, 1% NP-40, 1% sodium deoxycholate, 0.1% SDS, Life Technologies) containing fresh protease and phosphatase inhibitors (Life Technologies). Collected samples sat on ice for an additional 30 minutes and were pulled through a 28G insulin syringe for complete lysis. Protein concentrations were then determined using the Pierce BCA Protein Assay Kit (ThermoFisher Scientific). Equal amounts of protein samples (5-20µg) were diluted in RIPA buffer and βME-Laemmli buffer (Bio-Rad Laboratories) to equal volumes and boiled for 7 minutes at 95°C. Denatured samples were then loaded and run on Criterion TGX (Tris-Glycine eXtended) precast gels (Bio-Rad) for ∼15 minutes at 80V, and then ∼45 minutes at 150V. Migrated proteins were transferred from the gels to PVDF membranes using the Trans-Blot Turbo Transfer System (Bio-Rad) and run settings of 2.5A, 25V, for 7 minutes. Equal loading and complete transfer were checked with Ponceau S (Sigma Aldrich) staining for 30 minutes, shaking. After removing Ponceau, membranes were blocked for 45 minutes with shaking in 5% nonfat milk diluted in 1x TBS-T or SuperBlock T20 (Life Technologies) for phosphoproteins. Blocked membranes were incubated with primary antibodies overnight at 4°C, with shaking, using the following primary antibodies: Phospho-eIF2α (Cell Signaling Technology 9721S, 1:1000), eIF2α (Cell Signaling Technology 9722S, 1:1000), GRP78 BiP (Abcam ab21685, 1:1000), CHOP (Cell Signaling Technology 2895T, 1:1000), Cleaved Caspase-3 (Cell Signaling Technology 9664S, 1:1000), Actin (Cell Signaling Technology 3700S, 1:5000), β-Tubulin (Abcam ab6046, 1:10,000), GAPDH (Life Technologies AM4300, 1:5000).

Primary antibody solutions were removed after overnight incubation, and membranes washed 3 times with 1x TBS-T for 5 minutes. Membranes were then incubated for 1 hour, shaking, with species-matched secondary antibodies conjugated to horse radish peroxidase (Goat anti-Rabbit IgG (H+L) HRP, Life Technologies 31460, 1:5,000; Goat anti-Mouse IgG (H+L) HRP, Life Technologies 31430, 1:5000), diluted in block (5% milk or SuperBlock). Following 3 TBS-T washes, chemiluminescent signal was produced using the SuperSignal West Dura Extended Duration Substrate (ThermoFisher Scientific) and membrane signal was detected on film. All films were scanned using an EPSON scanner without automatic intensity contrast adjustments. Scanned images were cropped and measured for pixel intensity using FIJI/ImageJ. Figure quantifications display fold changes normalized for background film intensity and loading control protein.

### 4.7 RNA isolation, reverse transcription, and quantitative PCR

Total RNA was isolated from 2×10^6^ cells with Trizol (Life Technologies) according to manufacturer’s instructions. Briefly, cells were washed once with 1x PBS, on ice, and lysed with 250µL Trizol Reagent and mechanical disruption using a cell lifter. 50µL of chloroform was added to the Trizol-cell extract, and samples were centrifuged at 12,000g at 4°C for 15 minutes. The aqueous phase was collected into a clean Eppendorf tube and 125µL isopropanol, 15µg GlycoBlue Coprecipitant (Life Technologies) were added. Samples were incubated for 10 minutes at room temperature, and then centrifuged at 12,000g at 4°C for 10 minutes. The RNA pellet was washed twice with 250µL 75% ethanol, air-dried, and resuspended in 30µL of 1x DNase1 buffer (Life Technologies). 1µL of amplification grade DNase 1 was then added and incubated for 15 minutes at room temperature to degrade contaminating genomic DNA. 1µL of 25mM EDTA was added to each sample and incubated at 65°C for 10 minutes to quench the reaction.

Total RNA concentrations were analyzed using a nanodrop, and equal amounts of sample RNA were used to synthesize cDNA using the iScript Reverse Transcription Supermix for RT-qPCR (Bio-Rad). Per reaction, 4µL of 5X iScript RT supermix was combined with 0.5-1µg of total RNA, with water to a final volume of 20µL. Sample reactions were then run through the following thermocycler conditions- 5 minutes at 25°C, 20 minutes at 46°C, and 1 minute at 95°C.

Quantitative PCR (qPCR) was then performed on cDNA samples using Fast SYBR Green (Life Technologies). Per reaction, 5µL of 2X Fast SYBR Green Master Mix was combined with 1µL (50ng) of cDNA sample, 1µL of 10µM forward and reverse primer solution, and 3µL water. The following primer pairs were designed for target transcripts using Primer3Plus and validated using NCBI Blast and Geneious software (GAPDH forward 5’-TGACTTCAACAGCGACACCCA-3’, GAPDH reverse 5’-CACCCTGTTGCTGTAGCCAAA-3’; BiP forward 5’-TCTTCAGGAGCAAATGTCTTTGT-3’; BiP reverse 5’-CATCAAGTTCTTGCCGTTCA-3’; CHOP forward 5’-AGGGCTAACATTCTTACCTCTTCA-3’, CHOP reverse 5’-GATGAAAATGGGGGTACCTATG-3’). Technical triplicate reactions were set up for each sample and target primer in MicroAmp Optical 384-well Reaction Plates (Life Technologies), and reactions were run and analyzed on a Quant Studio 12K Flex Real-Time PCR system (ThermoFisher Scientific) following the standard Fast SYBER Green protocol. Melt curves for each target primer pair were analyzed to ensure amplification and quantification of a single PCR product. Differential gene expression was then determined using the ΔΔCT method and visualized as fold change values.

### 4.8 XBP1 splicing analysis

For XBP1 splicing analysis, 2µl (100ng) of cDNA samples were combined with 12.5µl Phusion Hot Start Flex 2X Master Mix (New England Biolabs), 2.5µL of 10µM forward and reverse primer solution (forward: 5’-GGGGCTTGGTATATATGTGG-3’, reverse: 5’-CCTTGTAGTTGAGAACCAGG-3’) and water to a final reaction volume of 25µl. Reactions were run with the following thermocycler conditions: 1 cycle of 98°C for 30 seconds; 40 cycles of 98°C for 10 seconds, 60°C for 30 seconds, and 72°C for 30 seconds; and 1 cycle of 72°C for 5 minutes. 2µg of resulting PCR products were then digested with Pst1-HF restriction enzyme (New England Biolabs) to cleave the unspliced XBP1 template. Digested PCR reactions were then run on a 2% agarose gel containing ethidium bromide at 135V. Bands were observed using a ChemiDoc XRS+ (Bio-Rad).

### 4.9 TUNEL assay

The Click-iT Plus TUNEL Assay for In Situ Apoptosis Detection using an Alexa Fluor 647 dye (Life Technologies) was performed according to manufacturer’s instructions. Cells were first fixed for 15 minutes at room temperature using 4% PFA diluted in 1x PBS. Samples were then permeabilized with 0.25% Triton X-100 in 1x PBS for 20 minutes at room temperature and washed twice with deionized water. Terminal deoxynucleotidyl transferase (TdT) reaction buffer was added to cells for 10 minutes at 37°C. TdT reaction buffer supplemented with TdT enzyme and EdUTP were then added for 60 minutes at 37°C for labeling of double-stranded DNA breaks. Following this incubation, cells were washed twice with 3% BSA in 1x PBS for 5 minutes each, and the Click-iT plus TUNEL reaction cocktail was added to cells at 37°C protected from light. Following 30 minutes of florescent labeling of EdUTP with click chemistry, the reaction mixture was removed, and cells were washed twice for 5 minutes with 3% BSA in 1x PBS. Cells were then stained with Hoechst, and immunofluorescent Isl1/2, and TUJ1 antibodies as normal for image acquisition and analysis of TUNEL+ MNs.

### 4.10 ER stress/protection assay and sample collection for proteomic and phosphoproteomic analyses

3×10^6^ patient derived MNs were plated in complete MN media on borate/poly-ornithine/laminin/fibronectin coated 6-well tissue culture plates. 3-day old cultures were treated with 0.1µM thapsigargin, 3µM kenpaullone, 5µM MAP4K4 inhibitor 29 or 0.1µM thapsigargin with 3µM kenpaullone or 5µM MAP4K4 inhibitor 29. 24 hours and 48 hours after compound treatment, MN cultures were washed 3 times with chilled PBS, on ice, and total protein was collected using 200µL/well of freshly prepared lysis buffer containing 6M urea (Life Technologies), 50mM EPPS (Sigma Aldrich), 1% triton X-100 (Sigma Aldrich), 5mM tris(2-carboxyethyl)phosphine (ThermoFisher Scientific), 20mM chloroacetamide (Sigma Aldrich), with 1x protease and phosphatase inhibitors (Life Technologies). Mechanical disruption with a cell lifter was used to collect the protein lysates into Eppendorf tubes which were then flash frozen in liquid nitrogen and stored at -80°C until mass spectrometry analysis.

### 4.11 Proteomics - Sample preparation and digestion

Protein concentrations from sample lysates were determined using the Bradford assay (ThermoFisher Scientific). Proteins denatured in 1% SDS were subjected to disulfide bond reduction with 5mM tris (2-carboxyethyl) phosphine (room temperature, 15 minutes) and alkylation with 20mM chloroacetamide (room temperature, 20 minutes). Methanol-chloroform precipitation was then preformed, adding 400μL of 100% methanol to the 100μg/100μL protein sample, vortexing 5 seconds, and then adding 100μL of 100% chloroform and vortexing 5 seconds. 300μL of water was added to the sample, vortexed 5 seconds, and centrifuged for 1 minute at 14,000g to generate distinct phase separations. The aqueous phase and organic phases were removed, leaving behind the protein disk that was washed twice with 400μL 100% methanol, and centrifuged at 21,000g for 2 minutes at room temperature. containing 0.1% RapiGest and digested at 37°C for 2 h with LysC protease at a 200:1 protein-to-protease ratio. Trypsin was then added at a 100:1 protein-to-protease ratio and the reaction was incubated for 6 h at 37°C.

Tandem mass tag labeling of each sample was performed by adding 10 µl of the 20 ng/μL stock of TMT reagent along with acetonitrile to achieve a final acetonitrile concentration of approximately 30% (v/v). Following incubation at room temperature for 1 h, labeling efficiency of a small aliquot was tested, and the reaction was then quenched with hydroxylamine to a final concentration of 0.5% (v/v) for 15 min. The TMT-labeled samples were pooled together at a 1:1 ratio. The sample was vacuum centrifuged to near dryness, resuspended in 5% formic acid for 15 min, centrifuged at 10000×g for 5 minutes at room temperature and subjected to C18 solid-phase extraction (SPE) (Sep-Pak, Waters).

### 4.12 Proteomics – TMT-labeled phosphopeptide enrichment

Phosphopeptides were enriched using Pierce Fe-NTA phosphopeptide enrichment kit (Thermo Fisher Scientific, A32992) following the provided protocol. In brief, combined TMT-labeled dried peptides were enriched for phosphopeptides, while the unbound peptides (flow through) and washes were combined and saved for total proteome analysis. The enriched phosphopeptides were dried down and fractionated according to manufacturer’s instructions using High pH reversed-phase peptide fractionation kit (Thermo Fisher Scientific, 84868) for a final 6 fractions and subjected to C18 StageTip desalting prior to MS analysis.

### 4.13 Proteomics - Off-line basic pH reversed-phase (BPRP) fractionation

Unbound TMT-labeled peptides (flow through from phosphopeptide enrichment step) and washes were dried down and resuspended in 100 μl of 10 mM NH4HCO3 pH 8.0 and fractionated using BPRP HPLC (Paulo et al. 2016). Briefly, samples were offline fractionated over a 90 min run, into 96 fractions by high pH reverse-phase HPLC (Agilent LC1260) through an aeris peptide xb-c18 column (Phenomenex; 250 mm x 3.6 mm) with mobile phase A containing 5% acetonitrile and 10 mM NH_4_HCO_3_ in LC-MS grade H_2_O, and mobile phase B containing 90% acetonitrile and 10 mM NH_4_HCO_3_ in LC-MS grade H_2_O (both pH 8.0). The 96 resulting fractions were then pooled in a non-continuous manner into 24 fractions (as outlined in Supplemental Fig. 5 of Paulo et al., 2016 (Paulo et al. 2016) and 12 fractions (even numbers) were used for subsequent mass spectrometry analysis. Fractions were vacuum centrifuged to near dryness. Each consolidated fraction was desalted via StageTip, dried again via vacuum centrifugation, and reconstituted in 5% acetonitrile, 1% formic acid for LC-MS/MS processing.

### 4.14 Proteomics - Liquid chromatography and tandem mass spectrometry

Mass spectrometry data were collected using an Orbitrap Fusion Lumos mass spectrometer (Thermo Fisher Scientific, San Jose, CA) coupled to a Proxeon EASY-nLC1200 liquid chromatography (LC) pump (Thermo Fisher Scientific). Peptides were separated on a 100 μm inner diameter microcapillary column packed in house with ∼35 cm of Accucore150 resin (2.6 μm, 150 Å, ThermoFisher Scientific, San Jose, CA) with a gradient consisting of 5%–16% (0-78 min), 16-22% (78-98min), 22-28% (98-110 min) (ACN, 0.1% FA) over a total 120 min at ∼500 nL/min. For analysis, we loaded 1/2 of each fraction onto the column. Each analysis used the Multi-Notch MS3-based TMT method (McAlister et al. 2014). The scan sequence began with an MS1 spectrum (Orbitrap analysis; resolution 120,000 at 200 Th; mass range 400−1400 m/z; automatic gain control (AGC) target 1×10^6^; maximum injection time 50 ms). Precursors for MS2 analysis were selected using a Top10 method. MS2 analysis consisted of collision-induced dissociation (quadrupole ion trap analysis; Turbo scan rate; AGC 2.0×10^4^; isolation window 0.7 Th; normalized collision energy (NCE) 35; maximum injection time 150 ms) with MultiStage Activation (MSA) for neutral loss of 97.9763. Monoisotopic peak assignment was used, and previously interrogated precursors were excluded using a dynamic window (150 s ±7 ppm). Following acquisition of each MS2 spectrum, a synchronous-precursor-selection (SPS) MS3 scan was collected on the top 10 most intense ions in the MS2 spectrum (McAlister et al., 2014). MS3 precursors were fragmented by high energy collision-induced dissociation (HCD) and analyzed using the Orbitrap (NCE 65; AGC 1.5×10^5^; maximum injection time 250 ms, resolution was 50,000 at 200 Th).

Mass spectrometry data were collected using an Orbitrap Fusion Lumos mass spectrometer (Thermo Fisher Scientific, San Jose, CA) coupled to a Proxeon EASY-nLC1200 liquid chromatography (LC) pump (Thermo Fisher Scientific). Peptides were separated on a 100 μm inner diameter microcapillary column packed in house with ∼35 cm of Accucore150 resin (2.6 μm, 150 Å, Thermo Fisher Scientific, San Jose, CA) with a gradient consisting of 5%–22% (0-125 min), 22-28% (125-140min) (ACN, 0.1% FA) over a total 150 min run at ∼500 nL/min. For analysis, we loaded 1/10 of each fraction onto the column. Each analysis used the Multi-Notch MS3-based TMT method (McAlister et al. 2014), to reduce ion interference compared to MS2 quantification. The scan sequence began with an MS1 spectrum (Orbitrap analysis; resolution 120,000 at 200 Th; mass range 350−1400 m/z; automatic gain control (AGC) target 5×10^5^; maximum injection time 50 ms). Precursors for MS2 analysis were selected using a Top10 method. MS2 analysis consisted of collision-induced dissociation (quadrupole ion trap analysis; Turbo scan rate; AGC 2.0×10^4^; isolation window 0.7 Th; normalized collision energy (NCE) 35; maximum injection time 35 ms). Monoisotopic peak assignment was used, and previously interrogated precursors were excluded using a dynamic window (150 s ±7 ppm), and dependent scan was performed on a single charge state per precursor. Following acquisition of each MS2 spectrum, a synchronous-precursor-selection (SPS) MS3 scan was collected on the top 10 most intense ions in the MS2 spectrum (McAlister et al. 2014). MS3 precursors were fragmented by high energy collision-induced dissociation (HCD) and analyzed using the Orbitrap (NCE 65; AGC 3×10^5^; maximum injection time 150 ms, resolution was 50,000 at 200 Th).

### 4.15 Proteomics - Data analysis

Mass spectra were processed using a Sequest-based (v.28, rev. 12) in-house software pipeline (Huttlin et al. 2010). Spectra were converted to mzXML using a modified version of ReAdW.exe. Database searching included all entries from the UniProt Human Reference Proteome database (2017 - SwissProt and TrEMBL). Sequences of common contaminant proteins (for example, trypsin, keratins and so on) were appended and the database was concatenated with one composed of all size-sorted protein sequences in reverse order. Searches were performed using a mass tolerance of 20 p.p.m. for precursors and a fragment-ion tolerance of 0.9 Da, and a maximum of two missed cleavages per peptide was allowed. TMT tags on lysine residues and peptide N termini (+229.163 Da) and carbamidomethylation of cysteine residues (+57.021 Da) were set as static modifications (except when testing for labeling efficiency, in which case the TMT modifications are set to variable), while oxidation of methionine residues (+15.995 Da) was set as a variable modification. Peptide-spectrum matches (PSMs) were adjusted to a 1% false discovery rate (FDR) and PSM filtering was performed using a linear discriminant analysis, as described previously (Huttlin et al. 2010), while considering the following parameters: Xcorr and Diff Seq. Delta Score, missed cleavages, peptide length, charge state, and precursor mass accuracy. Using the Picked FDR method (Savitski et al. 2015), proteins were filtered to the target 1% FDR level. Moreover, protein assembly was guided by principles of parsimony to produce the smallest set of proteins necessary to account for all observed peptides. For TMT-based reporter ion quantitation, we extracted the summed signal-to-noise (S:N) ratio for each TMT channel and found the closest matching centroid to the expected mass of the TMT reporter ion (integration tolerance of 0.003 Da). Proteins were quantified by summing reporter ion counts across all matching PSMs using in-house software, as described previously (Huttlin et al. 2010). PSMs with poor quality, MS3 spectra with more than 8 TMT reporter ion channels missing, or isolation specificity less than 0.7 (or 0.6 for phosphorylation dataset), or with TMT reporter summed signal-to-noise ratio that were less than 150 (100 for phosphorylation dataset) or had no MS3 spectra were excluded from quantification.

For phosphorylation dataset search, phosphorylation (+79.966 Da) on Serine, Threonine or Tyrosine and deamidation (+0.984 Da) on Asparagine or Glutamine were set as additional variable modifications. Phosphorylation site localization was determined using the AScore algorithm (Beausoleil et al. 2006). AScore is a probability-based approach for high-throughput protein phosphorylation site localization. Specifically, a threshold of 13 corresponded to 95% confidence in site localization.

Protein quantification values were exported for further analysis in Microsoft Excel and Perseus (Tyanova et al. 2016) and statistical test and parameters used are indicated in the corresponding Supplementary Data Set 1. Briefly, Welch’s t-test analysis was performed to compare two datasets, using s0 parameter (a minimal fold change cut-off) and correction for multiple comparison was achieved by the permutation-based FDR method, both functions that are built-in in Perseus software. For whole cell proteome (Figure 4A) analysis, each reporter ion channel was summed across all quantified proteins and normalized assuming equal protein loading of all samples. For phosphopeptide dataset (Figure 4C), peptide abundance was normalized to the protein abundance when available.

### 4.16 Gene ontology analyses

Gene ontology enrichment analyses were performed using the Database for Annotation, Visualization, and Integrated Discovery (DAVID) v6.8. Differentially expressed genes that reached a false discovery rate q-value significance threshold <0.05 were analyzed for enrichment compared to the total protein list quantified using default stringency parameters. GOTERM_BP_DIRECT terms with corresponding -log_10_(p-values) from the functional annotation chart results were graphed.

### 4.17 Kinase-substrate enrichment analyses

Kinase-substrate enrichment analyses were performed on Supplementary Data Set 1 using the KSEA software, available as the R package ‘KSEAapp’ on CRAN: CRAN.R-project.org/package=KSEAapp/ and online at https://casecpb.shinyapps.io/ksea/ (Wiredja, Koyutürk, and Chance 2017). The log_2_(fold change) of phosphoproteins in each treatment-comparison group were used as input, and kinase-substrate annotations were derived from both PhosphoSitePlus and NetworKIN datasets. The resulting output displayed a z-score for each kinase, which described the collective phosphorylation status of its substrates, such that a kinase with a negative score had substrates that were generally dephosphorylated with the test group, and vice versa for a kinase with a positive score. P values for kinase z-scores were then determined by assessing the one-tailed probability of having a more extreme score than the one measured, followed by a Benjamini-Hochberg false discovery rate correction for multiple hypothesis testing. Kinase z-scores with p<0.05 were determined significant, and unbiased hierarchical clustering was performed to determine the relative similarity of kinase scores between each treatment group.

### 4.18 DNA extraction and genotyping PCR

Genomic DNA was extracted using the Wizard Genomic DNA Purification Kit (Promega). Approximately 2×10^6^ cells were lysed using nuclei lysis solution and mechanical trituration by pipetting. RNase solution was added to the nuclear lysate and incubated for 15 minutes at 37°C. Protein Precipitation Solution was added, vortexed briefly, and centrifuged for 4 minutes at 16,000g. Sample solutions were mixed gently by inversion until white thread-like strands of DNA were visible. Samples were centrifuged for 1 minute at 16,000g, and the DNA pellet was washed with 70% ethanol before centrifuging again (1 minute, 16,000g). The DNA pellet was air dried for 15 minutes and resuspended with DNA rehydration solution for 1 hour at 65°C or overnight at 4°C. Total DNA concentrations were analyzed using a nanodrop.

Polymerase chain reaction (PCR) was performed using the Phusion Hot Start Flex 2X Master Mix (New England Biolabs). Reactions were set up according to the manufacturer’s instructions. (TDP-43^G298S^ forward 5’-CGACTGAAATATCACTGCTGCTG-3’, TDP-43^G298S^ reverse 5’-GGATGCTGATCCCCAACCAA-3’; SOD1^L144F^ forward 5’- GTTATTTTTCTAATATTATGAGG- 3’, SOD1^L144F^ reverse 5’- GTTTTATAAAACTATACAAATCTTCC-3’; SOD1A4V forward 5’- GTTTGGGGCCAGAGTGGG-3’, SOD1A4V reverse 5’-CCGGGCGCTGGACCAGGGCGGCCCC-3’). Thermocycler conditions were the following- 1 cycle of 98°C for 30 seconds; 30 cycles of 98°C for 10 seconds, 45-72°C for 30 seconds (depending on melting temperature of primer pairs determined by NEB Tm calculator https://tmcalculator.neb.com), 72°C for 30 seconds/kb; and 1 cycle of 72°C for 10 minutes. PCR products were run at 135V on a 1% agarose gel containing ethidium bromide and bands were observed using a ChemiDoc XRS+ (Bio-Rad). Sequencing of PCR products was performed by Psomagen and resulting sequencing files were analyzed with Geneious software using NCBI gene data.

### 4.19 Statistical Analyses

All statistical analyses were performed using rstatix: Pipe-Friendly Framework for Basic Statistical Tests, R package version 0.6.0. https://CRAN.R-project.org/package=rstatix (Alboukadel Kassambara, 2020). Data were determined to be normally distributed using the Shapiro Wilk test and variance was determined equal between groups using Levene’s test. For comparisons of 2 groups, a 2 tailed, unpaired students t-test was used. For comparisons of 3 or more groups, an analysis of variance (ANOVA) was used followed by Tukey’s HSD post hoc tests. p<0.05 was considered statistically significant and denoted in graphs with a *, p<0.01 **, p<0.001 ***, and P<0.0001 ****.

## 5 Conflict of Interest

J.W.H. is a founder and consultant for Caraway Therapeutics and a founding scientific advisor for Interline Therapeutics. L.L.R. is a founder of Elevian, Rejuveron, and Vesalius Therapeutics, a member of their scientific advisory boards and a private equity shareholder. All are interested in formulating approaches intended to treat diseases of the nervous system and other tissues. He is also on the advisory board of Alkahest, a Grifols company, focused on the plasma proteome, and scientific advisory board member of ProjenX and Corsalex. None of these companies provided any financial support for the work in this paper. However, both ProjenX and Corsalex are focused on treating ALS, including via modulating targets that are described in this paper.

## 6 Author Contributions

M.E.W. and L.L.R. conceived and designed the study; M.E.W. performed iPSC culture, MN differentiation and culture, ER stressor and protection assays, IF staining, image acquisition and analysis, western blot and qRT-PCR analyses, and gene ontology analyses; A.O. performed proteomics and phosphoproteomic analyses under supervision by J.W.H., K.M.H performed gene set enrichment analyses, kinase-substrate enrichment analyses and statistics. M.E.W. wrote the manuscript; R.M.G. edited the manuscript and performed revision experiments. All authors reviewed and approved manuscript submission.

## 7 Funding

This work was supported by Biogen, Harvard Brain Initiative ALS Grant, and gifts from Alkahest, Inc., and Google to L.L.R. This work was also supported by the NIH (R37 NS083524, R01 NS110395 to J.W.H. and F32 AG079593 to R.M.G.).

## 8 Acknowledgments

We thank Dr. Chen Wu and Dr. Silvia Piccinotti for helpful suggestions and discussions. Figure 6 was created with BioRender.com.

