## Supplementary Data and Material for "Analyzing ER stress response in ALS patient derived motor neurons identifies druggable neuroprotective targets": Supplemental Figures.pdf

### Figure S1. Extended analyses of iPSC-derived MN cultures and response to proteostatic stressors.

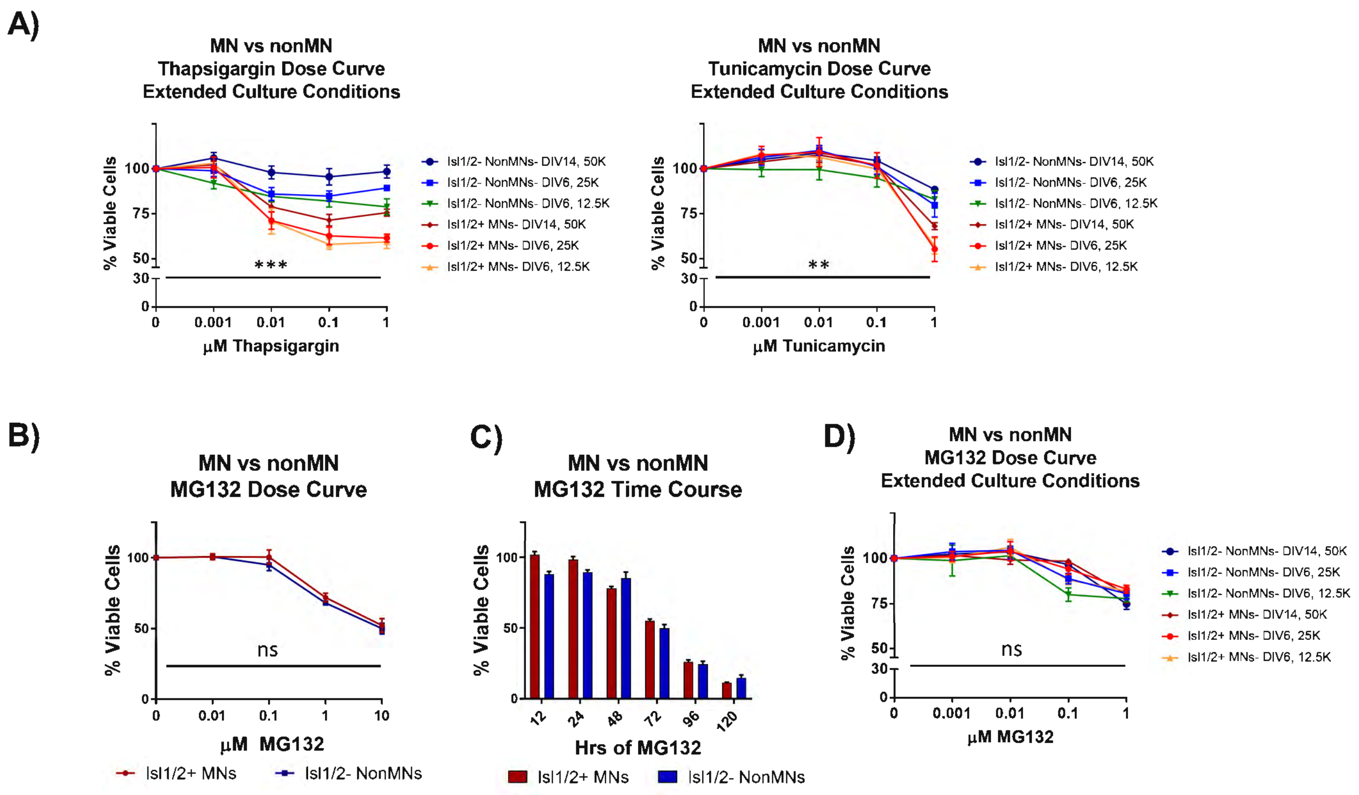

### Figure S2. Overview of global phosphoproteomics experiment.

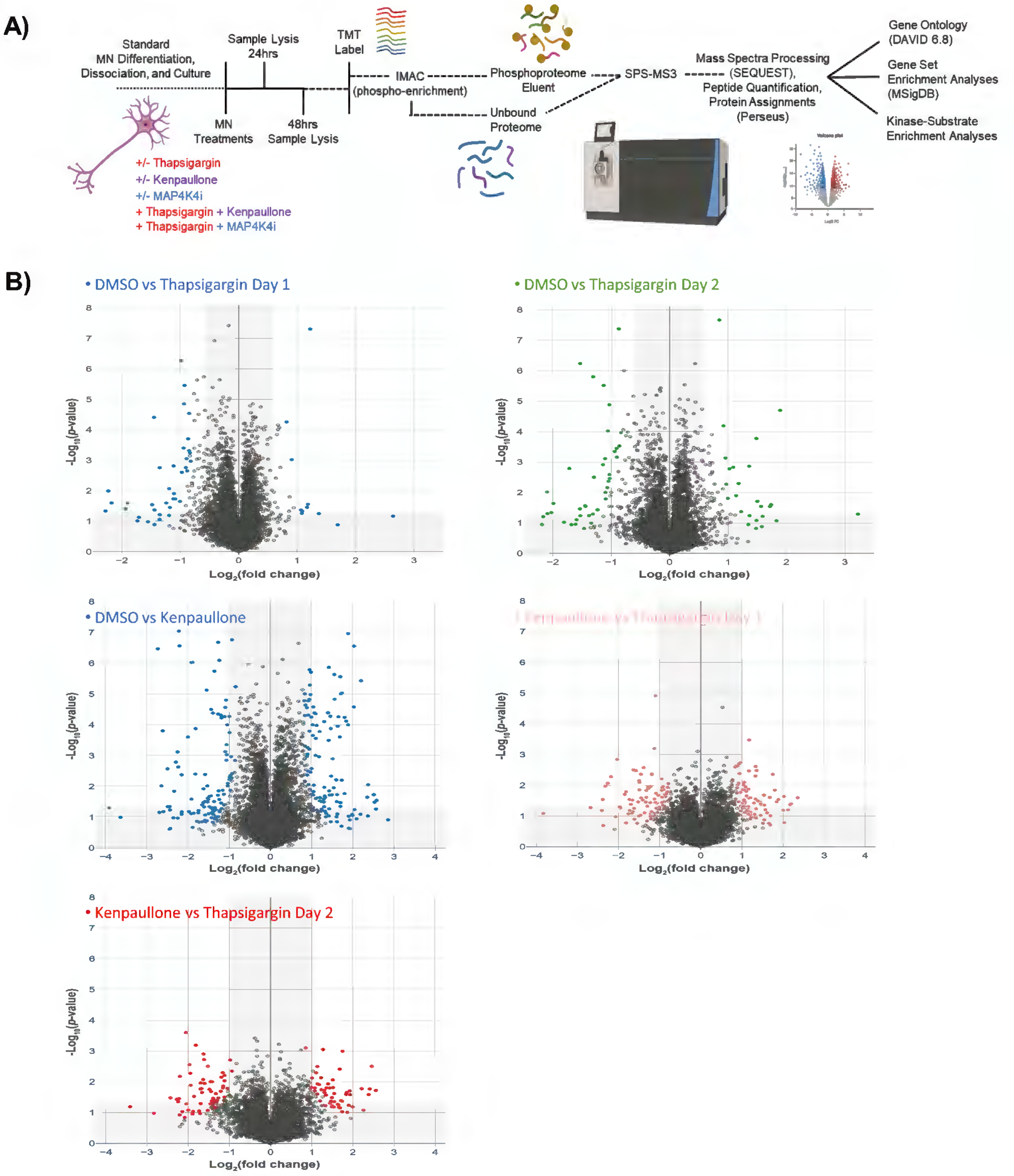

#### Figure S3. Approach to identifying viable MNs.

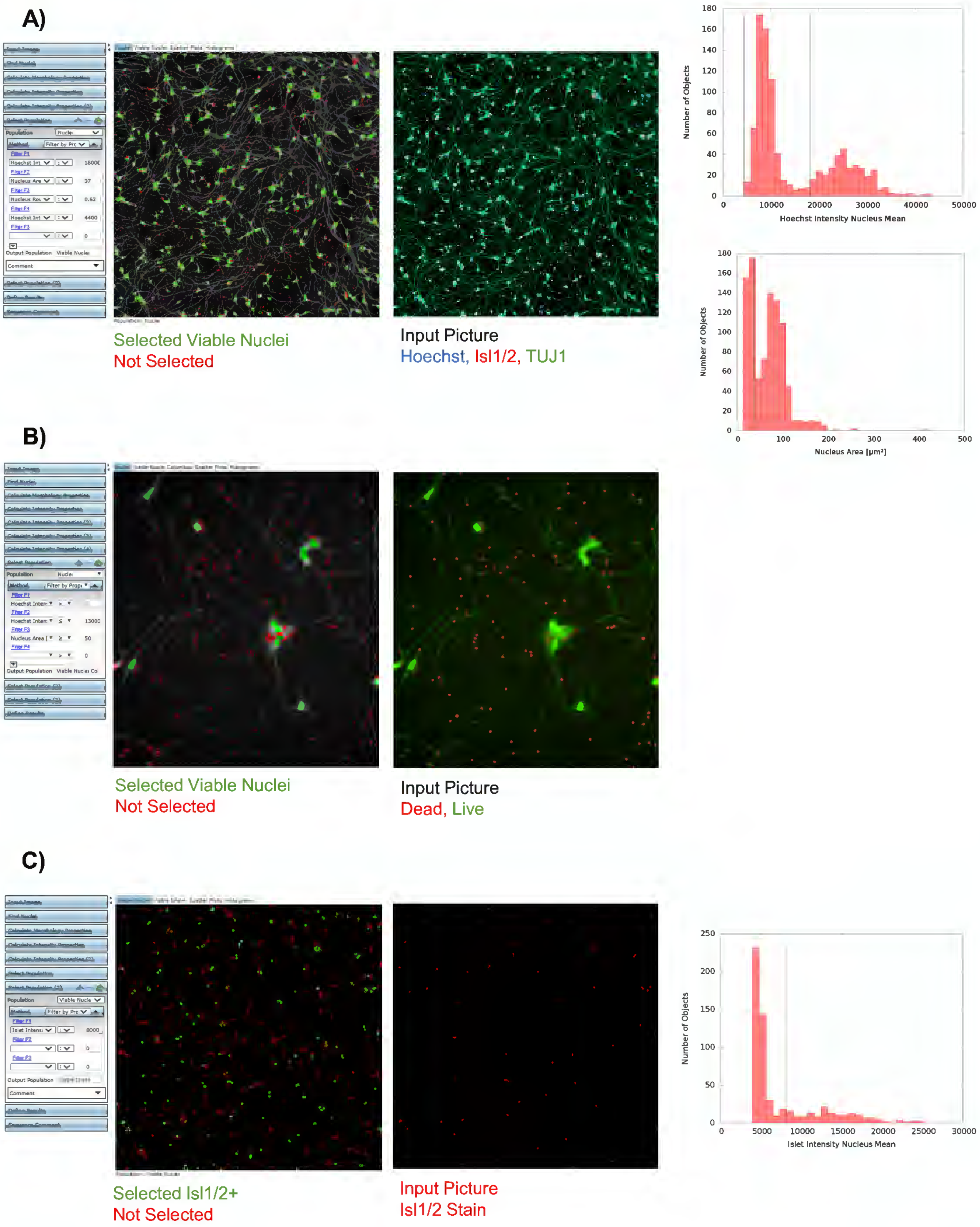

### Figure S4. Neurite tracing of β-Tubulin III staining.

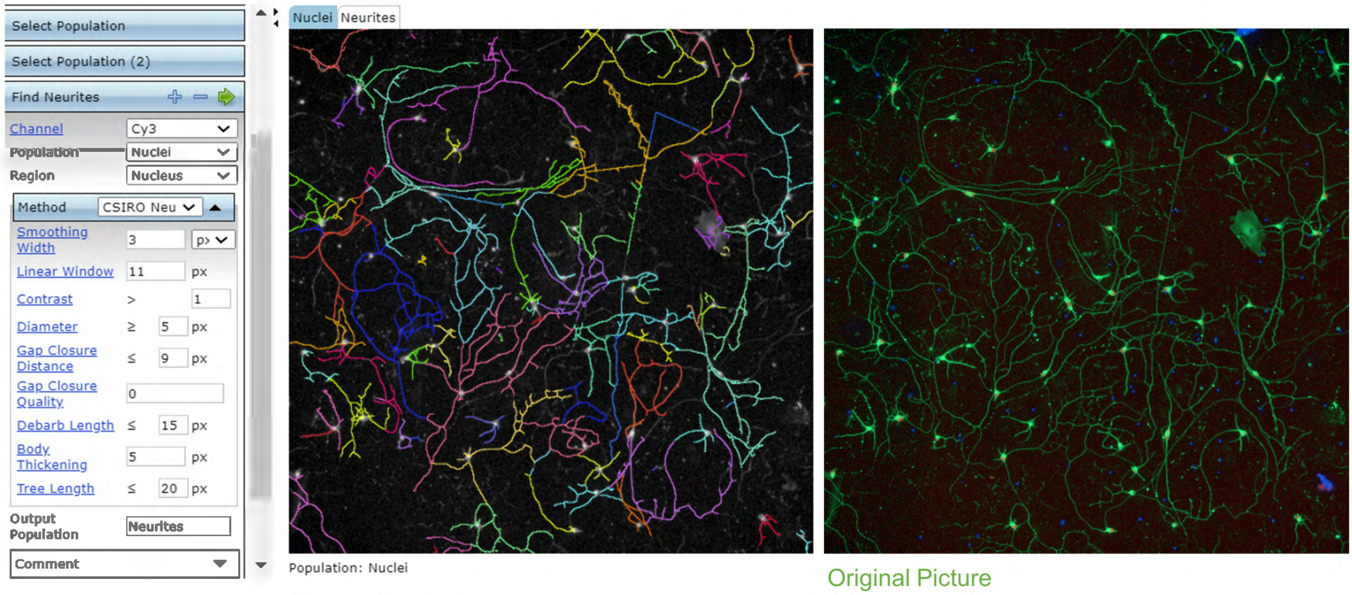


Neurites Traced

Original Picture
β-Tubulin III Stain
