## Supplementary Data and Material for "Analyzing ER stress response in ALS patient derived motor neurons identifies druggable neuroprotective targets": Supplementary Material_20231021.docx

**Supplementary Figure S1. Extended analyses of iPSC-derived MN cultures and response to proteostatic stressors.** (A) MN and non-MN viability from low density cultures (25K/96 well or 12.5K/96 well) or mature cultures (14 days in vitro (DIV) at 50K/96well) after 48hrs exposure to increasing doses of ER stressors. Nb=3, nt=2, two-way ANOVA; 25K thapsigargin- p = 7.67x10^-6^, 12.5K thapsigargin- p = 0.000258, 2week thapsigargin- p = 0.000691, 25K tunicamycin- p = 0.008. 12.5K tunicamycin- p = 2.63x10^-7^, 2week tunicamycin- p = 0.002. For simplicity, stars indicating significance are shown for the condition with the least significance. (B) Quantification of MN and non-MN viability 48hrs after treatment with increasing concentrations of MG132. Nb=3, nt=12, two-way ANOVA, p = 0.394. (C) Quantification of MN and non-MN viability after treatment with 1μM MG132 for various lengths of time. (D) MN and non-MN viability from low density cultures (25K/96 well or 12.5K/96 well) or mature cultures (14 days in vitro (DIV) at 50K/96well) after 48hrs exposure to increasing doses of MG132. Nb=3, nt=2, two-way ANOVA; 25K- p = 0.894. 12.5K- p = 0.947. 2week- p = 0.284. Biological replicate experiments denoted as Nb, each with technical replicate experiments nt. Data are mean value +/- SEM.

**Supplementary Figure S2. Overview of global phosphoproteomics experiment.** (A) Schematic of the ER stress and protection assay and the subsequent quantitative proteomics analysis pipeline. (B) Individual, separated volcano plots from Figure 4A and C.

**Supplementary Figure S3. Approach to identifying viable MNs.** (A) Example nuclear size exclusion parameters and Hoechst intensity thresholding used to identify the viable cell population. Histograms to right of selection script and input image demonstrate 2 distinct cell populations, live or dead, with live cells demonstrating a nuclear area >~37-55μm2 and Hoechst intensities lower than the threshold brightness of pyknotic nuclei (18,000 in this example). (B) Viable cell script accuracy confirmed with LIVE/DEAD Viability/Cytotoxicity Kit, for mammalian cells (Life Technologies L3224). (C) Example Isl1/2 intensity thresholding used to identify the viable MN population. Histogram to right of selection script and input image demonstrate that selected Isl1/2+ cell populations must have an Isl1/2 intensity greater than the basal intensity (>8000 in this example).

**Supplementary Figure S4. Neurite tracing of β-Tubulin III staining**. Representative image analysis pipeline to track neurites using B-Tubulin III staining.

**Supplementary Data Set 1. ER stress and protection proteomics and phosphoproteomics dataset.** Quantified proteins and phosphoproteins are displayed with corresponding log_2_foldchange with each treatment, compared to indicated control.

**Supplementary Video Files**. Automated live cell imaging of iPSC-derived MN cultures treated with DMSO (1), 1uM thapsigargin (2), or 1uM tunicamycin (3). Images were taken every 6hrs for 48hrs.

**Supplementary Table S1. Overview of quantified phosphopeptides and peptides**

| **Data Set** | **# Peptides (Set 1)** | **# Peptides (Set 2)** | **# Unique Proteins** | **# Unique Phosphorylation Sites** |
| --- | --- | --- | --- | --- |
| Stress+Protection (Protein) | 67,357 | 59,708 | 6,697 |  |
| Stress+Protection (Phospho-protein) | 28,399 | 28,973 | 2,764 | 7,999 |
